# Intracellular Calcium links Milk Stasis to Lysosome Dependent Cell Death by Activating a TGFβ3/TFEB/STAT3 Pathway Early during Mammary Gland Involution

**DOI:** 10.1101/2022.05.17.492309

**Authors:** Jaekwang Jeong, Jongwon Lee, Gabriel Talaia, Wonnam Kim, Junho Song, Juhyeon Hong, Kwangmin Yoo, David G. Gonzalez, Diana Athonvarangkul, Jaehun Shin, Pamela Dann, Ann M Haberman, Lark Kyun Kim, Shawn M. Ferguson, Jungmin Choi, John Wysolmerski

## Abstract

Involution of the mammary gland after lactation is a dramatic example of coordinated cell death. Weaning causes distension of the alveolar structures due to the accumulation of milk, which, in turn, activates STAT3 and initiates a caspase-independent but lysosome-dependent cell death (LDCD) pathway. Although the importance of STAT3 and LDCD in early mammary involution is well established, it has not been entirely clear how milk stasis activates STAT3. In this report, we demonstrate that protein levels of the PMCA2 calcium pump are significantly downregulated within 2- 4 hours of experimental milk stasis. Reductions in PMCA2 expression correlate with an increase in cytoplasmic calcium *in vivo* as measured by multiphoton intravital imaging of GCaMP6f fluorescence. These events occur concomitant with the appearance of nuclear pSTAT3 expression but prior to significant activation of LDCD or its previously implicated mediators such as LIF, IL6 and TGFβ3, all of which appear to be upregulated by increased intracellular calcium. We also observed that milk stasis, loss of PMCA2 expression and increased intracellular calcium levels activate TFEB, an important regulator of lysosome biogenesis. This is the result of increased TGFβ signaling and inhibition of cell cycle progression. Finally, we demonstrate that increased intracellular calcium activates STAT3 by inducing degradation of its negative regulator, SOCS3, a process which also appears to be mediated by TGFβ signaling. In summary, these data suggest that intracellular calcium serves as an important proximal biochemical signal linking milk stasis to STAT3 activation, increased lysosomal biogenesis, and lysosome-mediated cell death.

## Introduction

Involution of the mammary gland following lactation is one of the most dramatic examples of coordinated cell death in nature (1–4). This process is initiated by the failure to empty milk from the gland for more than 12-24 hours, which results in distension of the alveolar structures, a change in the shape of mammary epithelial cells (MECs) and programmed cell death of many epithelial cells. This first phase of involution is regulated by local mechanisms and is reversible. If the gland remains unsuckled for more than 48-72 hours, a second phase of irreversible involution ensues, characterized by widespread cell death, proteolytic disruption of the basement membranes, and remodeling of epithelial and stromal components of the gland to approximate its pre-pregnant structure (1–4).

Although several pathways have been implicated in triggering the initial phase of involution (1, 4–12), a principal mediator of this process appears to be the activation of signal transducer and activator 3 (STAT3), which has been thought to occur due to the secretion of cytokines, such as leukemia inhibitory factor (LIF), interleukin-6 (IL6) and transforming growth factor (TGF) β3 by MECs in response to milk stasis (8-10, 13, 14). STAT3, in turn, increases the number and size of lysosomes in MECs as well as the expression of lysosomal enzymes such as cathepsin B and L (8, 15, 16). Together, these events result in a caspase-independent form of lysosome-dependent cell death (LDCD) (8, 16). Interestingly, a similar process of LDCD occurs in neurons in response to ischemia-reperfusion injury, where it is triggered, in part, by cellular calcium (Ca^2+^) overload (17, 18).

MECs transport large amounts of Ca^2+^ from the systemic circulation into milk, a process involving the plasma membrane calcium-ATPase 2 (PMCA2) (19). PMCA2 is expressed in the apical plasma membrane of MECs specifically during lactation (19, 20) and it transports Ca^2+^ out of cells in response to ATP hydrolysis (21–23). In its absence, milk Ca^2+^ transport is reduced by 60-70% (19, 20). PMCA2 levels decline rapidly after weaning and PMCA2-null mice demonstrate inappropriate, widespread MEC death during lactation (20). Given that PMCA2 is important for Ca^2+^ secretion from MECs, we hypothesized that the decline in PMCA2 levels upon weaning triggers LDCD by increasing intracellular Ca^2+^ levels. We now report that decreased PMCA2 expression is associated with increased cytoplasmic Ca^2+^ levels in MECs *in vivo*, and that increased intracellular Ca^2+^ triggers LIF, IL-6 and TGFβ3 expression, as well as STAT3 phosphorylation. Furthermore, we demonstrate that elevated intracellular Ca^2+^ levels activate transcriptional programs leading to lysosome biogenesis. These results suggest that an increase in intracellular Ca^2+^ due to reduced PMCA2-mediated calcium clearance represents an important proximal event coupling milk stasis to LDCD.

## Results

### Milk Stasis Increases Intracellular Calcium Levels

We hypothesized that a decrease in PMCA2 expression after weaning might increase intracellular Ca^2+^ levels within mammary epithelial cells and, as a result, contribute to lysosome-dependent cell death (LDCD). Experimentally, involution can be initiated in a single mouse mammary gland by sealing the teat with adhesive (24). Given that the other 9 glands are suckled normally and continue to make milk, this model isolates the consequences of milk stasis from systemic changes caused by weaning. As shown in Fig. 1A, we confirmed that, compared to the contralateral suckled gland (control), PMCA2 immunofluorescence was significantly reduced by 4 hours after teat sealing and was decreased to very low levels by 24 hours (20). This was associated with an increase in nuclear staining for pSTAT3, which was initially detectable at 4 hours after teat sealing and progressively increased at 8 and 24 hours (Fig. 1A&B).

**Figure 1.**
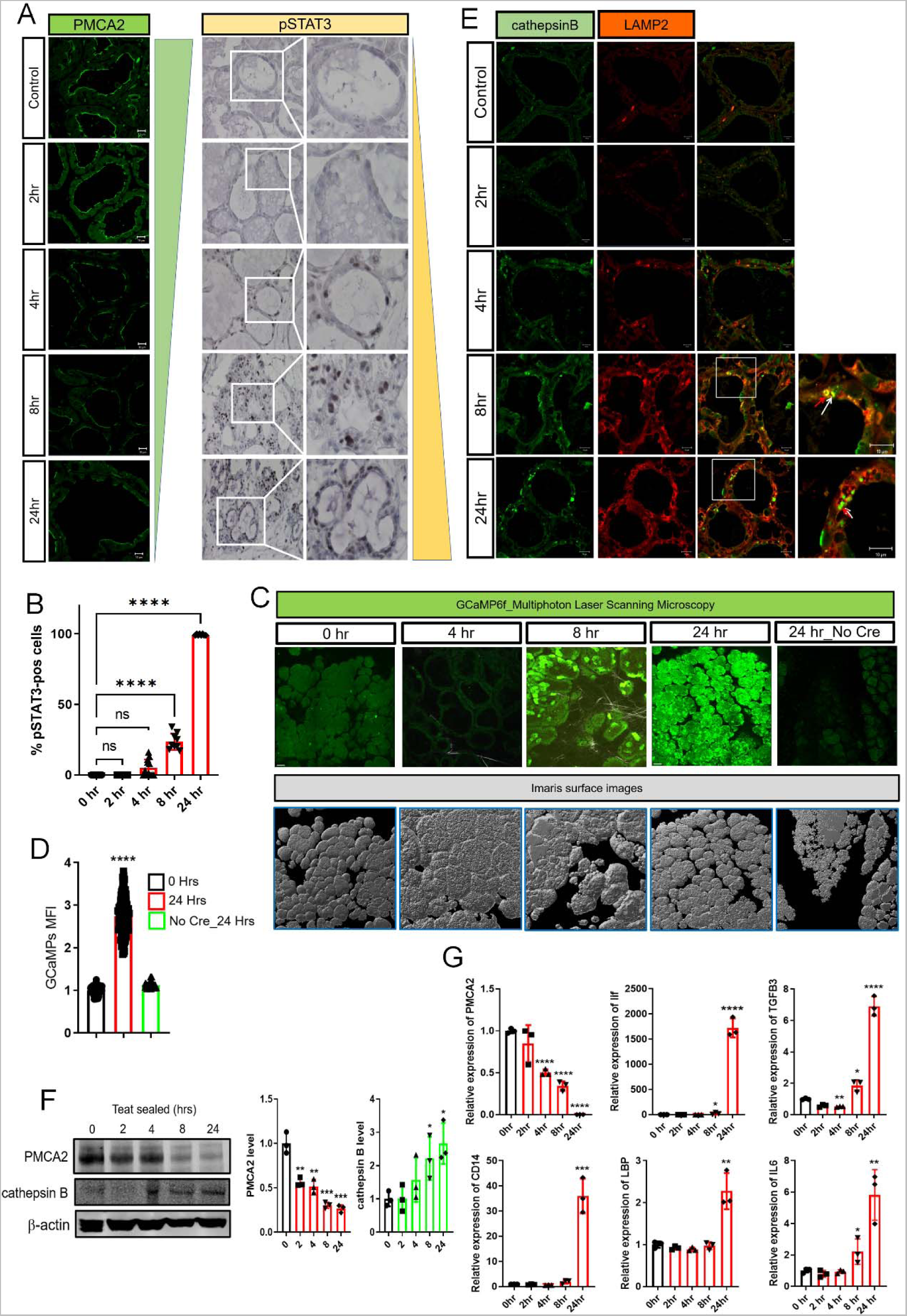
Milk Stasis Increases Intracellular Calcium Levels. A) Immunofluorescence for PMCA2 and Immunohistochemistry for pSTAT3 at 0, 2, 4, 8, and 24 hours post teat-sealing mammary gland. (representative images of n=3). 0 hours represents the unsealed contralateral lactating gland. B) Quantification of the percentage of pSTAT3-positive epithelial cells in Figure 1. C) Images obtained using multiphoton laser scanning microscopy at 0, 4, 8 and 24 hours post teat-sealing of a mammary gland using BLG-Cre;Ai95 females. The contralateral, unsealed gland served as a control (0 hours). Bottom row shows matching imaris surface images. (representative images of n=3). D) Mean Fluorescent Intensity (MFI) of MECs at 0 and 24 hours post teat-sealing (n=3). Mice without Cre expression served as controls. E) Immunofluorescence for Cathepsin B and LAMP2 at 0, 2, 4, 8, and 24 hours post teat- sealing mammary gland. (representative images of n=3). F) Western blot analysis of PMCA2 and Cathepsin B at 0, 2, 4, 8, and 24 hours post teat-sealing mammary gland. (n=3). G) PMCA2, Lif, TGFβ3, CD14, LBP, and IL6 mRNA expression in mammary glands at 0, 4, 8, and 24 hours post teat-sealing, as assessed by quantitative RT-PCR (QPCR) (n=3). All scale bars represent 10μm.

In order to test whether milk stasis increased cytoplasmic Ca^2+^ levels *in vivo*, we generated transgenic mice expressing the genetically encoded, GCaMP6f calcium sensor specifically in MECs. GCaMP6f is a modified EGFP, which fluoresces with progressive intensity in response to increasing concentrations of cytoplasmic Ca^2+^ (25, 26). We crossed Ai95(RCL-GCaMP6f)-D (Ai95) mice to mice expressing Cre recombinase under the control of the beta-lactoglobin promoter (BLG-Cre mice) to activate expression of GCaMP6f in MECs at the transition from pregnancy to lactation (27). Lactating BLG-Cre;Ai95 females underwent intravital imaging of mammary epithelial cells using multiphoton laser scanning microscopy of anesthetized mice at 4, 8 and 24 hours post teat-sealing of a 4^th^ inguinal mammary gland; the contralateral gland, which was not sealed, served as a control. We could not image individual mice sequentially for a full 24 hours. Therefore, mice were either studied at 4 and 8 hours, or at 24 hours after teat sealing. We used the second harmonic-generated signal from the collagen fibers within the fascia covering the glands as an anchoring point to ensure that we compared images at comparable tissue depths. As shown in Fig. 1C&D, and in Supplemental Video.1-4, teat-sealing caused a significant increase in intracellular Ca^2+^ levels *in vivo*, as evidenced by an increase in GCaMP fluorescence, which was first detectable at 8 hours and which persisted, increased, and became more uniform at 24 hours. There was no similar increase of fluorescence 24 hours after teat sealing in the glands of Ai95 females in the absence of Cre recombinase (Fig. 1C&D, Supplemental Video 5). These data demonstrate that milk stasis increases cytoplasmic Ca^2+^ levels in mammary epithelial cells *in vivo*.

The regulation of early involution is complex but factors implicated as important triggers of LDCD include leukemia inhibitory factor (LIF), interleukin 6 (IL6) and transforming growth factor-beta 3 (TGFβ3), all of which are produced by MECs in response to milk stasis and can contribute to STAT3 activation (8-10, 13, 14). In turn, STAT3 signaling has been shown to be required for LDCD in MECs after weaning (8, 28, 29). In order to assess how temporal changes in PMCA2 expression and cytoplasmic Ca^2+^ concentration correlate with the onset of LDCD, we compared them to changes in LAMP2, cathepsin B, LIF, IL6 and TGFβ3 expression at 2,4,8 and 24 hours post teat sealing. The contralateral gland which continued to be suckled normally served as a control in all experiments. During lactation, staining intensity for LAMP2 and cathepsin B was low but both progressively increased after teat sealing, beginning with the 4-hour time point. Immunofluorescence demonstrated an increase in the size of defined lysosomes that co-stained for both LAMP2 and Cathepsin B but also an increase in more diffuse cytoplasmic staining for both markers (Fig. 1E). At 24 hours, we observed the appearance of intensely staining foci of Cathepsin B that did not co- localize with LAMP2 staining. These changes were mirrored by increasing levels of Cathepsin B as assessed by immunoblotting whereas PMCA2 levels declined during this same time course (Fig. 1F). Interestingly, we did not see any change in LIF mRNA expression until 8 hours and this increase was minor compared to the prominent increase in LIF mRNA at 24 hours post teat sealing (Fig. 1G). A similar pattern was seen for TGFβ3 and IL6 mRNA expression, and we actually noted a decrease in TGFβ3 mRNA expression at 2 and 4 hours post teat sealing, before observing an increase in its expression over baseline at 8 and 24 hours. The relative increases in IL6 and TGFβ3 mRNA levels were quantitatively much less than the increase in LIF mRNA. We also examined the expression of two STAT3-target genes that have been noted to participate in the inflammatory responses to involution, LBP (lipopolysaccharide binding protein) and CD14 (Lipopolysaccharide receptor) (7, 12). Both mRNAs were only significantly elevated after 24-hours of teat sealing. These changes all occurred either after or concurrent with the decrease in PMCA2 expression or the increase in intracellular Ca^2+^ levels, but not before. Therefore, decreased PMCA2 expression is an early response to milk stasis, occurring before significant increases in cytoplasmic Ca^2+^ concentrations, widespread STAT3 activation or upregulation of LDCD markers. Interestingly, these data also demonstrate that decreased PMCA2 and increased pSTAT3 occur prior to significant increases in LIF, IL6 or TGFβ3 mRNA expression.

### Changes in PMCA2 and pSTAT3 expression are reversible with reintroduction of suckling

The first phase of mammary gland involution is reversible if pups are reintroduced to suckle within 48-hours of milk stasis (1, 3, 4, 7). We next examined whether the reversal of early involution would be associated with changes in PMCA2 expression and/or intracellular calcium. As illustrated in Fig. 2A, we compared mammary glands from 3 groups of lactating mice: A) mice who were sacrificed on day 10 of lactation without manipulation; B) mice whose pups were removed for 24 hours before the mothers were sacrificed; and C) mice whose pups were removed for 24 hours and then replaced to re-suckle for 24 hours before the mothers were sacrificed. As expected from the previous teat-sealing experiments, PMCA2 mRNA and protein levels were significantly reduced at 24 hours after pup withdrawal (Fig. 2B-D). Remarkably, 24 hours after pups were re-introduced, PMCA2 mRNA levels had recovered almost to baseline lactating levels (Fig. 2B). In addition, PMCA2 protein levels and apical membrane PMCA2 staining intensity were both increased back towards the levels noted during lactation (Fig. 2C&D). These changes in PMCA2 expression were associated with reciprocal changes in cytoplasmic Ca^2+^ as assessed by GCaMP6f fluorescence (Fig. 2E, Supplemental Video 6-7). Intracellular Ca^2+^ levels increased with weaning but were reduced back to baseline after suckling was reestablished. Like teat sealing, pup withdrawal also led to an increase in LIF, TGFβ3 and IL6 mRNA expression, but re-suckling restored the expression of all 3 genes back to lactating levels (Fig. 2F). Likewise, nuclear pSTAT3 staining as well as CD14 and LBP mRNA levels were increased by pup withdrawal but were suppressed back to baseline after reintroduction of the pups (Fig. 2F&G). Interestingly, nuclear staining for pSTAT5 persisted for 24-hours of pup withdrawal, at which time MECs expressed both pSTAT5 and pSTAT3 in their nuclei.

**Figure 2.**
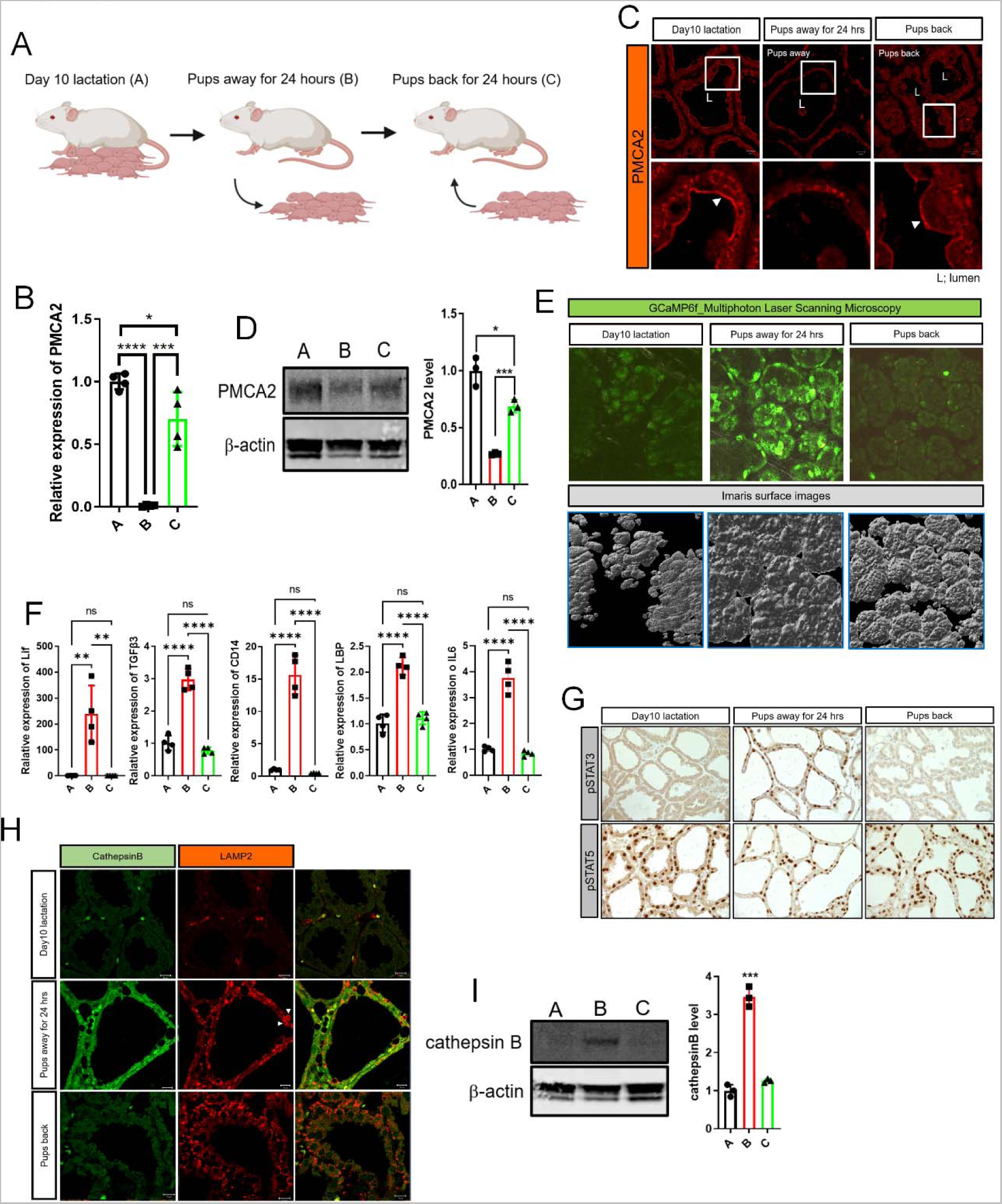
Changes in PMCA2, intracellular calcium and pSTAT3 levels are reversible with reintroduction of suckling. A) Experimental design. A, day 10 lactation as a control; B, 24 hours after pup removal at day 10 of lactation, C; 24 hours after pup reintroduction following 24-hours without suckling. Created with BioRender.com. B) PMCA2 mRNA expression, assessed by QPCR from conditions A, B, and C (n=4). C) Immunofluorescence for PMCA2 from tissue sections of mammary glands from A, B, and C. White arrows; apical plasma membrane. (n=4). D) Western blot analysis of PMCA2 from mammary tissue extracts from A, B, and C. (n=4). E) Representative Images obtained using multiphoton laser scanning microscopy of mammary glands from conditions A, B, and C using BLG-Cre;Ai95 females. Bottom row shows imaris surface images. (n=3). F) lif, TGFβ3, CD14, LBP, and IL6 mRNA levels in mammary glands harvested from conditions A, B, and C, assessed by QPCR (n=4). G) Immunohistochemistry for pSTAT3 and pSTAT5 in mammary glands harvested from conditions A, B, and C. H) Immunofluorescence for Cathepsin B and LAMP2 in mammary glands harvested from conditions A, B, and C. White arrows; enlarged lysosomes. I) Western blot analysis of Cathepsin B from tissue extracts of mammary glands harvested from conditions A, B, and C. (n=4). All scale bars represent 10μm.

We also performed immunofluorescence staining and immunoblotting to assess cathepsin B expression. As seen in Figs. 2H&I, cathepsin B levels were increased at 24-hours of pup withdrawal and, like pSTAT3 staining, Cathepsin B fluorescence and protein levels were reduced back towards baseline 24 hours after reintroduction of the pups. In contrast, LAMP2 immunofluorescence intensity was increased by pup withdrawal but remained elevated 24-hours after pups were reintroduced (Fig. 2H). These data demonstrate strong correlations between PMCA2 expression, intracellular Ca^2+^ levels, activation of STAT3 and induction of LDCD markers.

### Loss of PMCA2 prematurely activates LDCD during lactation

We next examined mediators of LDCD in PMCA2-null mice in order to determine whether loss of PMCA2 was sufficient to activate this pathway. As noted previously, loss of PMCA2 expression caused premature activation of STAT3 (20) as evidenced by pSTAT3 expression in the nuclei of MECs at mid-lactation and an increase in pSTAT3 levels on immunoblots of whole mammary glands (Supplemental Fig. 1A&B). Mammary glands from lactating PMCA2-null mice also demonstrated an increase in LAMP2 and Cathepsin B immunofluorescence as well as an increase in cathepsin B levels (Supplemental Fig. 1C&D). Significantly, loss of PMCA2 caused an increase in LIF, TGFβ3 and IL6 mRNA levels during mid-lactation, even though continued suckling by pups removed milk, preventing milk stasis and alveolar distension (Supplemental Fig. 1E). As in the teat-sealed and weaned glands (Figs. 1&2) expression of the STAT3 targets, CD14 and LBP was also upregulated inappropriately during lactation (Supplemental Fig. 1E). These data demonstrate that loss of PMCA2 is sufficient to prematurely induce cytokine expression, STAT3 activation and LDCD in MECs during lactation, suggesting that the early decline in PMCA2 levels in response to milk stasis, serves as an important trigger for LDCD during involution.

### Intracellular Calcium activates STAT3 by degrading SOCS3 in a TGFβ-dependent pathway

In order to test whether loss of PMCA2 triggers LDCD by raising intracellular Ca^2+^, we treated MCF10A cells, an immortalized but non-transformed human mammary epithelial cell line, with ionomycin and increased extracellular Ca^2+^ to increase cytoplasmic Ca^2+^ levels (30). We examined changes in cytoplasmic Ca^2+^ by measuring the increase in fluorescence in MCF10A cells transiently transfected with a Ca^2+^ indicator, RCaMP (31). As expected, treating MCF10A cells with 1μM ionomycin and 10mM extracellular Ca^2+^ for 16 hours uniformly increased RCaMP fluorescence, resembling the pattern of GCaMP6f fluorescence at 24 hrs after teat sealing (Supplemental Fig 2A and Fig. 1C). It also increased nuclear pSTAT3 expression, lysosomal mass and immunofluorescence for cathepsin B and LAMP2 (Supplemental Fig 2B-D). We also observed an induction of LIF, IL6 and TGFβ3 mRNA expression as well as the STAT3 target genes, CD14 and LBP (Supplemental Fig. 2F). These data demonstrate that, in MCF10A cells *in vitro*, increased levels of intracellular Ca^2+^ are sufficient to trigger key aspects of the STAT3-LDCD pathway.

In MECs, STAT3 stimulates *Suppressor of Cytokine Signaling 3 (SOCS3)* gene expression and, in turn, SOCS3 inhibits STAT3 phosphorylation (32–34), defining a short negative feedback loop (Fig. 3A). Moreover, similar to the findings in PMCA2-null mammary glands, deletion of SOCS3 from MECs causes inappropriate activation of STAT3 during lactation (34). Therefore, we next examined SOCS3 levels in response to teat sealing and in PMCA2-null mammary glands. As shown in Fig. 3B, as compared to the contralateral lactating gland (time 0), SOCS3 protein levels were diminished within 2-4 hours after teat sealing and became substantially reduced by 8 and 24 hours. Interestingly, this occurred despite a marked increase in *Socs3* mRNA levels at 8 and 24-hours after teat sealing (Fig. 3C). SOCS3 levels were also significantly reduced in lactating PMCA2-null glands as compared to wild-type lactating glands (Fig. 3D). As with teat-sealing, the pattern was the opposite for *Socs3* mRNA levels; despite the decrease in SOCS3 protein levels, there was a significant increase in *Socs3* mRNA levels in lactating PMCA2-null mammary glands as compared to wild-type control lactating glands (Fig. 3E). Similarly, reintroduction of suckling after 24-hours of weaning led to reciprocal changes in SOCS3 protein and mRNA levels. Re-suckling increased SOCS3 protein (Fig. 3F) and reduced *Socs3* mRNA levels (Fig. 3G).

**Figure 3.**
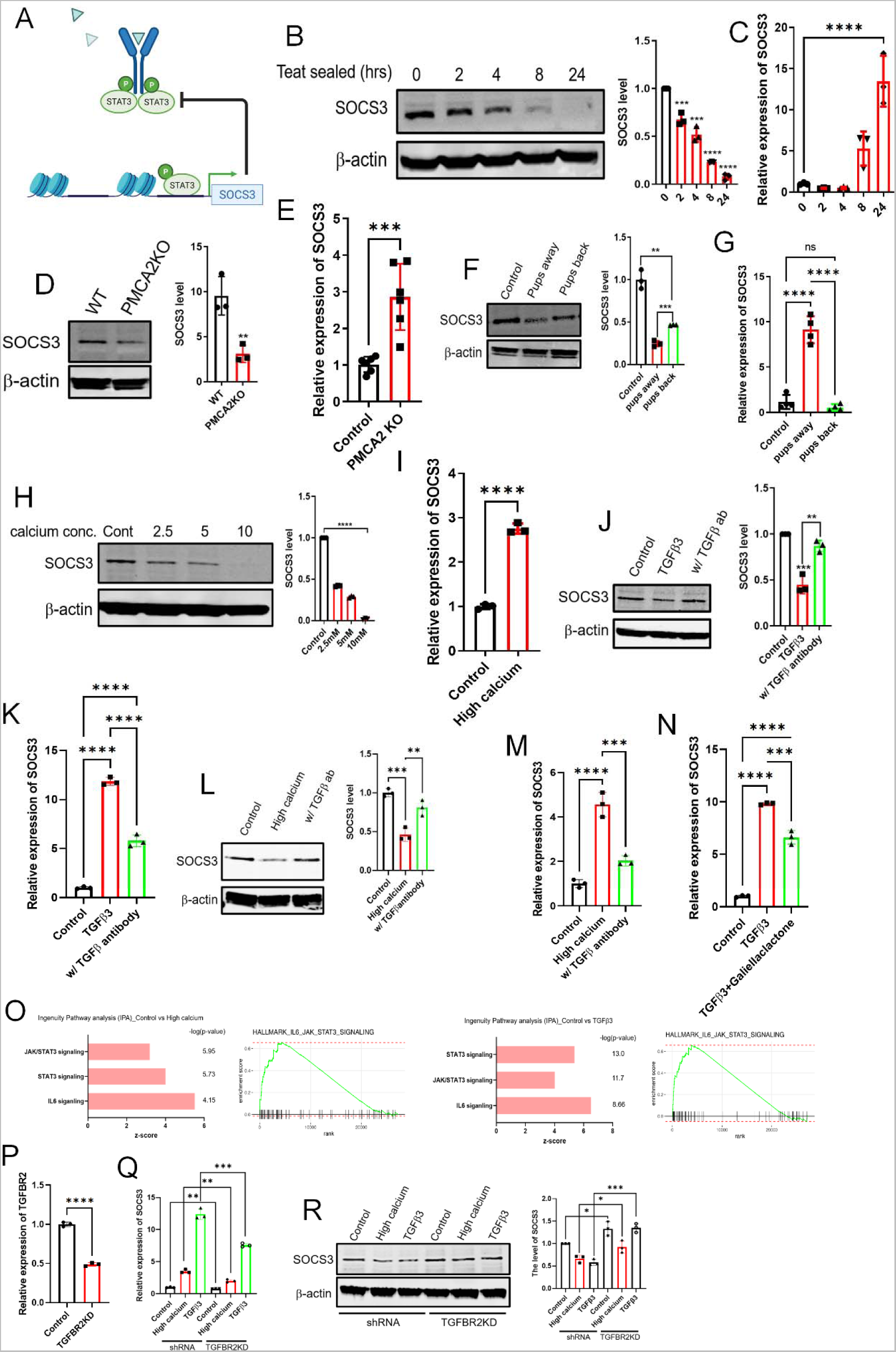
Calcium mediated TGFβ signaling activates STAT3 signaling by degrading SOCS3 protein. A) Diagram showing negative feedback of SOCS3 on STAT3 signaling. Created with BioRender.com. B) Western blot analysis of SOCS3 in mammary glands harvested at 0, 2, 4, 8, and 24 hours post teat-sealing. (n=3). C) Socs3 mRNA expression assessed by QPCR in mammary glands harvested at 0, 2, 4, 8, and 24 hours post teat-sealing (n=3). D) Western blot analysis of SOCS3 in mammary glands from control and PMCA2 KO mice harvested on day 10 of lactation. (n=3). E) SOCS3 mRNA expression assessed by QPCR in mammary glands from control and PMCA2 KO mice harvested on day 10 of lactation. (n=6). F-G) Western blot and QPCR analysis of SOCS3 from tissue extracts harvested from control lactating mice, mice 24 hrs. after teat sealing, and mice with re-suckling of mammary glands for 24 hours. (n=4). H) Western blot analysis of SOCS3 in MCF10A cell exposed to increasing concentrations of extracellular calcium (0, 2.5, 5, 10mM) + 1μM ionomycin. (n=3). I) SOCS3 mRNA expression assessed by QPCR in MCF10A cells exposed to control and high calcium (10mM calcium + 1μM ionomycin) conditions (n=3). J-K) Western blot and QPCR analysis of SOCS3 in MCF10A cells exposed to 10ng/ml TGFβ3 ± 2μg/ml pan-TGFβ antibody. (n=3). L) Western blot analysis of SOCS3 levels in MCF10A cells exposed to high calcium conditions ± 2μg/ml pan-TGFβ antibody. (n=3). M) SOCS3 mRNA expression in MCF10A cells under high calcium condition ± pan- TGFβ antibody, assessed by QPCR. (n=3). N) SOCS3 mRNA expression assessed by QPCR in MCF10A cells exposed to TGFβ3 ± 10μM/ml of Galiellaclactone (n=3). O) Ingenuity Pathways Analysis (IPA) from RNAseq compared between control MCF10A cells versus either high calcium or TGFβ3 which highlights the activation of JAK-STAT3 signaling. P) TGFBR2 mRNA expression in control and TGFBR2 knock down MCF10A cells. (n=3). Q) SOCS3 mRNA expression assessed by QPCR in control and TGFBR2 knock down MCF10A cells exposed to high calcium and TGFβ3 (n=3). R) Western blot analysis of SOCS3 in control and TGFBR2 knock down MCF10A cells exposed to high calcium and TGFβ3 (n=3).

Given that *Socs3* gene expression is induced by STAT3, increased *Socs3* mRNA levels likely reflect the fact that SOCS3 protein is degraded in response to loss of PMCA2 expression, which we hypothesized was caused by increased intracellular Ca^2+^ (35–37). To test this possibility, we examined the effects of increasing intracellular Ca^2+^ on SOCS3 protein and mRNA expression in MCF10A cells. As shown in Fig. 3H, treatment of these cells with ionomycin and increasing doses of extracellular Ca^2+^ resulted in a dose-dependent decrease in SOCS3 protein levels as assessed by immunoblot. Increasing intracellular Ca^2+^ levels also caused a reciprocal increase in Socs3 mRNA levels (Fig. 3I). Together, these data suggest that the loss of PMCA2 and the resulting increased levels of intracellular Ca^2+^ due to milk stasis, lead to SOCS degradation, which, in turn, increases pSTAT3 levels.

TGFβ and Jak-Stat signaling pathways interact in many settings (38, 39), and since TGFβ3 expression was induced in response to teat-sealing (Fig. 1G), loss of PMCA2 expression (Supplemental Fig. 1E) and increased intracellular Ca^2+^ levels in MCF10A cells (Supplemental Fig. 2E), we next asked whether TGFβ signaling contributed to the reductions in SOCS3 levels noted in MCF10A cells. As shown in Supplemental Fig. 3A, and has been reported previously, at 24-hours after teat sealing, the expression of both TGFβ1 and TGFβ3 is increased in the mammary gland, although TGFβ3 expression predominates, and TGFβ2 levels remain unchanged (1, 10). We next treated MCF10A cells with each of the three isoforms of TGFβ and measured SOCS3 protein levels and SOCS3 mRNA levels. As shown in Fig. 3J&K, as well as Supplemental Fig. 3B&C, TGFβ3 and TGFβ1 reduced SOCS3 protein levels but increased SOCS3 mRNA levels. TGFβ2 had no effect on either SOCS3 protein or mRNA levels. Of note, the effects of either TGFβ3 or TGFβ1 could be blunted using an antibody that is described to inhibit all TGF-beta isoforms (40, 41). Furthermore, treatment of MCF10A cells with the same TGFβ-blocking antibody was able to blunt the ability of 10mM calcium plus ionomycin to reduce SOCS3 protein levels (Fig. 3L) and to induce *SOCS3* mRNA expression (Fig. 3M). Finally, simultaneous treatment of cells with TGFβ3 or TGFβ1 and a STAT3 inhibitor, galiellaclatone, abrogated the rise in SOCS3 mRNA levels supporting the idea that the increase in SOCS3 mRNA expression in response to TGFβ was due to the dis-inhibition of STAT3 signaling secondary to the decline in SOCS3 protein (Fig. 3N, Supplemental Fig 3D).

In order to validate these findings, we treated MCF10A cells with either high calcium or with TGFβ3 as described above and performed bulk RNA sequencing. As shown in Fig. 3O, treatment of MCF10A cells with either calcium and ionomycin or with TGFββ3 significantly altered the expression of genes in the canonical JAK/STAT signaling pathway, in the canonical STAT3 pathway and in the canonical IL6 signaling pathway as determined by Ingenuity Pathway Analysis. Furthermore, gene set enrichment analysis (GSEA) also demonstrated upregulation of genes within the hallmark IL6-JAK-STAT3 signaling pathway. We next knocked down TGFBR2 (Transforming growth factor beta receptor II) in MCF10A cells (TGFBR2KD) (Fig. 3P). As expected, high calcium and TGFβ3 induced SOCS3 gene expression but reduced SOCS3 protein levels in control cells (Fig 3Q&R). In contrast, knocking down TGFBR2 blunted the effects of calcium or TGFβ3 on inducing SOCS3 mRNA levels (Fig. 3Q). SOCS3 protein levels were increased in the TGFBR2-knockdown cells at baseline and calcium and TGFβ3 were less effective at reducing SOCS3 protein levels in these cells as compared to controls (Fig. 3R). Together, these data combined with those presented in the prior paragraphs, suggest that the effects of intracellular Ca^2+^ levels on SOCS3 expression and STAT3 activation may be mediated, in part, by TGFβ signaling.

### Increased intracellular calcium levels activate TFEB signaling to increase lysosomal biogenesis

The initiation of LDCD involves an increase in lysosome mass as well as increases in lysosome membrane permeability (8, 16). Consistent with this observation, we found expression of the lysosomal marker, LAMP2, to be increased in response to teat sealing, pup withdrawal and in lactating PMCA2-null glands (Figs. 1&2, Supplemental Fig. 1 & Fig. 4A). An important regulator of lysosome biogenesis is the transcription factor EB (TFEB) (42, 43) and immunostaining for TFEB revealed increased total and nuclear staining in both the teat-sealed mammary gland and in lactating PMCA2-null glands (Fig. 4B). TFEB mRNA levels were also increased in response to teat sealing and in PMCA2-null glands during lactation (Fig. 4C). Likewise, withdrawal of pups for 24 hours increased TFEB mRNA levels, but reintroduction of pups after 24 hours of weaning, led to a reduction of TFEB mRNA levels back to below baseline levels, correlating with changes in PMCA2 expression and intracellular Ca^2+^ levels as noted previously (Fig. 4D, Fig.2). We also detected changes in gene expression consistent with activation of TFEB when we interrogated a RNAseq database which included mouse mammary glands harvested at day 10 of lactation and at day 2 of involution (44). KEGG-identified gene set enrichment analysis demonstrated an increase in the expression of genes associated with lysosomes and autophagy at day 2 of involution as compared to lactation, both of which are regulated by TFEB (Fig. 4E). Furthermore, as shown in the volcano plot and associated heat maps, involution was associated with an increase in the expression of genes previously identified as specifically regulated by TFEB, including multiple lysosomal proteins, such as the cathepsin and Lamp families (Fig. 4F&G).

**Figure 4.**
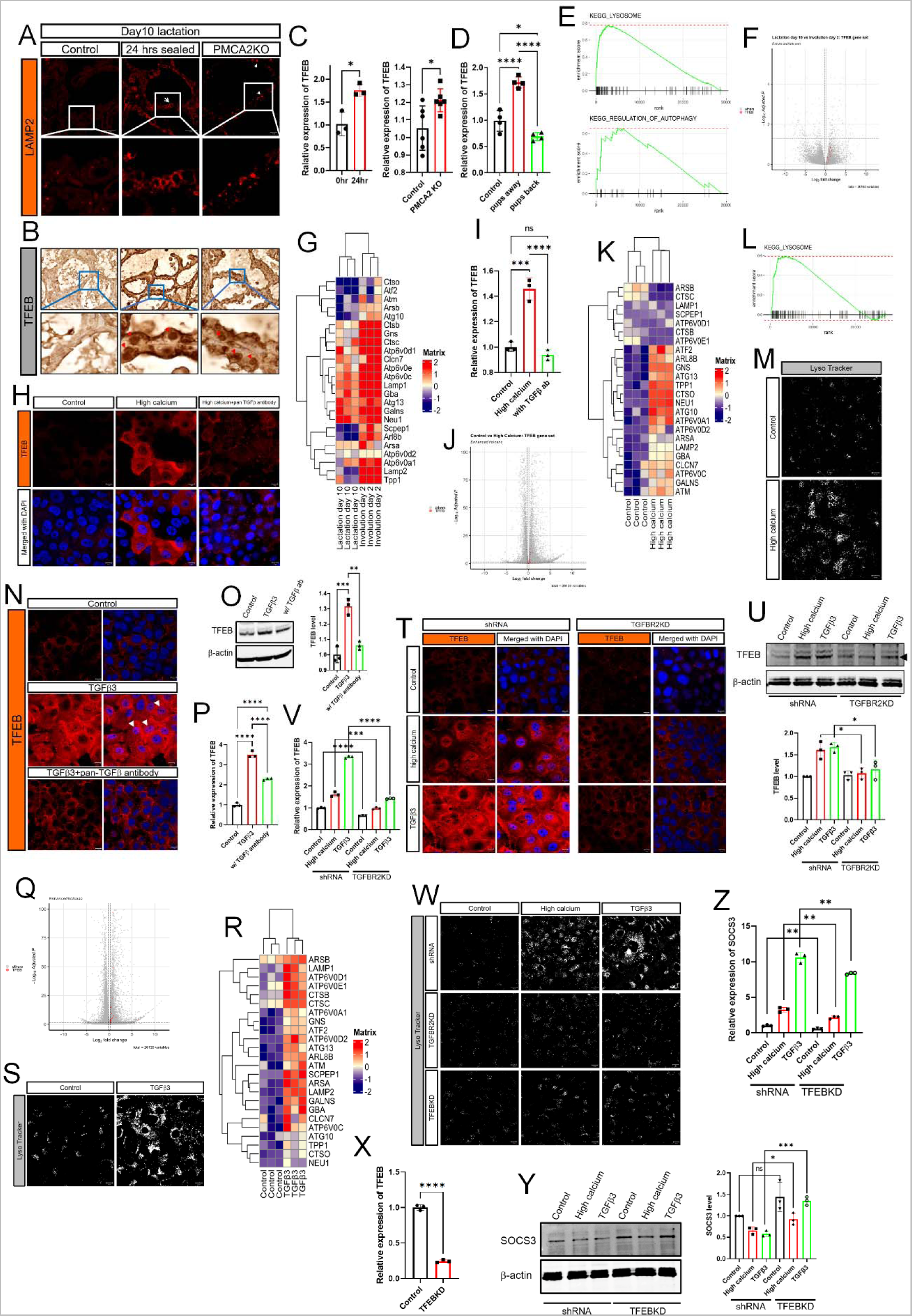
Intracellular Calcium and TGFβ increase TFEB and lysosomal biogenesis. A) Immunofluorescence for LAMP2 in mammary glands from control mice on day 10 of lactation, 24 hrs after teat-sealing, and PMCA2 KO mice on day 10 of lactation. B) Immunohistochemistry for TFEB in mammary glands from control mice on day 10 of lactation, 24 hrs after teat-sealing, and PMCA2 KO mice on day 10 of lactation. C) TFEB mRNA expression assessed by QPCR in mammary glands harvested from control mice on day 10 of lactation, 24 hrs after teat-sealing, and PMCA2 KO mice on day 10 of lactation (n=6). D) TFEB mRNA expression assessed by QPCR in mammary glands harvested from control lactating mice, mice 24 hrs. after teat sealing, and mice with re-suckling of mammary glands for 24 hours (n=4). E) KEGG gene set enrichment plot of lysosome and autophagy pathway from RNAseq results compared between day 10 lactation and day 2 involution in the mammary gland. F-G) Volcano and Heatmap plot which highlight TFEB-regulated genes from RNAseq results compared between day 10 lactation and day 2 involution in the mammary gland. H) Immunofluorescence for TFEB in MCF10A cells exposed to high calcium ± pan-TGFβ antibody. I) TFEB mRNA expression in MCF10A cells exposed to high calcium ± pan- TGFβ antibody, assessed by QPCR. (n=3). J-K) Volcano and Heatmap plots highlight TFEB-regulated genes from RNAseq results in MCF10A cells in control versus high calcium conditions. L) KEGG gene set enrichment plot of lysosome pathway from RNAseq results compared between control and high calcium condition in MCF10A cells. M) Live cell images stained by LysoTracker in MCF10A cells at control and high calcium conditions. N) Immunofluorescence for TFEB in MCF10A cells exposed to TGFβ3 ± pan-TGFβ antibody. O) Western blot analysis of TFEB in MCF10A cells exposed to TGFβ3 ± pan-TGFβ antibody. (n=3). P) TFEB mRNA expression assessed by QPCR in MCF10A cells exposed to TGFβ3± pan-TGFβ antibody. (n=3). Q-R) Volcano and Heatmap plots highlighting TFEB-regulated genes from RNAseq results comparing MCF10A cells at baseline and after treatment with TGFβ3. S) LysoTracker staining in MCF10A cells at baseline and after treatment with TGFβ3. All scale bars represent 10μm. T) Immunofluorescence for TFEB in control and TGFBR2 knock down MCF10A cells exposed to high calcium and TGFβ3. All scale bars represent 10μm. U) Western blot analysis of TFEB in control and TGFBR2 knock down MCF10A cells exposed to high calcium and TGFβ3. (n=3). V) TFEB mRNA expression assessed by QPCR in control and TGFBR2 knock down MCF10A cells exposed to high calcium and TGFβ3 (n=3). W) LysoTracker staining in control, TGFBR2-, and TFEB- knock down MCF10A cells exposed to high calcium and TGFβ3. All scale bars represent 10μm. X) TFEB mRNA expression in control and TFEB knock down MCF10A cells. (n=3). Y) Western blot analysis of SOCS3 in control and TFEB knock down MCF10A cells exposed to high calcium and TGFβ3 (n=3). Z) SOCS3 mRNA expression assessed by QPCR in control and TFEB knock down MCF10A cells exposed to high calcium and TGFβ3 (n=3).

These results *in vivo* were reinforced by results in mammary epithelial cells *in vitro*. As shown in Fig. 4H, incubating MCF10A cells with 1μM ionomycin and 10mM extracellular Ca^2+^ for 16 hours led to a dramatic increase in immunofluorescence for TFEB, both in the cytoplasm and in nuclei. This was associated with an increase in TFEB mRNA levels (Fig. 4I). In addition, analysis of RNAseq data from MCF10A cells treated with high calcium demonstrated activation of genes known to be regulated by TFEB and KEGG-identified gene set enrichment analysis demonstrated a significant increase in the expression of genes associated with lysosomes (Fig. 4J-L). LysoTracker staining confirmed the increase of lysosomal mass in MCF10A cells treated with high calcium (Fig. 4M). Given the results regarding SOCS3 levels, we also asked whether TFEB levels could be regulated by TGFβs in MCF10A cells. TGFβ1 and 3 reproduced the effects of high calcium by increasing total and nuclear immunofluorescence for TFEB (Fig. 4N, Supplemental Fig. 3E) as well as increasing overall TFEB protein by Western analysis (Fig. 4O, Supplemental Fig. 3F). TFEB mRNA levels were increased as well (Fig. 4P, Supplemental Fig. 3G). In each case, the induction of TFEB expression was mitigated by treatment with a pan-TGFβ blocking antibody (Fig. 4N-P). This antibody was also effective in blocking the induction of TFEB protein and mRNA levels by high calcium, suggesting that TGFβ3 mediates the effects of calcium in MCF10A cells (Fig. 4H&I). As with high calcium, analysis of RNAseq data from MCF10A cells treated with TGFβ3 demonstrated activation of a series of genes known to be regulated by TFEB (Fig. 4Q&R), and both TGFβ1 and TGFβ3 increased lysosomal mass (Fig. 4S, Supplemental Fig. 3H). We also analyzed the effects of high calcium and TGFβ3 treatment on TGFBR2 knockdown cells. As shown in Fig. 4T&U, knocking down TGFBR2 prevented the increases in TFEB immunofluorescence and total protein in response to treatment of control cells with either high calcium or TGFβ3. Similar results were noted for TFEB mRNA expression (Fig. 4V). Consistent with these results, knocking down TGFBR2 prevented the increase in lysosomal mass in response to high calcium and TGFβ3 treatment (Fig. 4W). In fact, inhibiting TGFβ signaling was as effective at knocking down TFEB itself in these cells (Fig. 4W&X). Finally, we used TFEB-knockdown MCF10A cells to determine whether the induction of TFEB might influence the ability of high calcium and TGFβ3 to affect SOCS3 levels. As shown in Figs 4Y&Z, knocking down TFEB resulted in higher baseline levels of SOCS3 protein and blunted the effects of both calcium and TGFβ3 on reducing SOCS3 levels. This resulted in a modest reduction in the ability of either calcium or TGFβ3 to induce SOCS3 gene expression, suggesting that TFEB affected SOCS3 but was not the sole mediator of the effects of calcium or TGFβ3.

### Activation of TFEB signaling by elevations in intracellular calcium or TGFβ3 is associated with inhibition of cell cycle progression

Nuclear translocation of TFEB is controlled by multiple mechanisms, including by calcineurin signaling, by mTOR signaling and by the cell-cycle regulators, CDK4/6 (45, 46). Increasing intracellular calcium levels in MCF10A cells activated the GCaMP3- TRPML1 calcium sensor, demonstrating an increase in lysosomal Ca^2+^ content and export (47) (Supplemental Fig. 4A). As expected, this was also associated with nuclear translocation of NFAT, a standard bioassay of calcineurin activity (48) (Supplemental Fig. 4B). However, TGFβ3 did not similarly activate NFAT, and treatment of MCF10A cells with the calcineurin inhibitor, cyclosporin A, did not prevent the increase in expression of 2 TFEB target genes, LAMP2 and Cathepsin B, in response to TGFβ3 (Supplemental Fig. 4B&C). We also examined whether treating MCF10A cells with calcium and ionomycin or TGFβ3 inhibited mTOR signaling, which has been reported to cause nuclear accumulation of TFEB (49, 50). However, treatment with calcium and ionomycin or with TGFβ3 appeared to promote mTOR activity (Supplemental Fig. 4D). Thus, neither activation of calcineurin activity nor inhibition of mTOR appeared to explain the increase in total and nuclear TFEB in response to calcium/ionomycin or TGFβ3.

We next focused on the possibility that alterations in cell cycle regulation might affect TFEB expression and/or nuclear translocation given that TGFβ signaling is known to modulate cell cycle progression (51) and that CDK4/6 have been shown to phosphorylate TFEB, inhibiting its nuclear localization (46). In support of this possibility, ingenuity pathway analysis of the previously described mammary gland RNAseq data comparing lactating and 48hrs of involution showed that involution was associated with upregulation of the “senescence” pathway and downregulation of the “cyclin and cell cycle regulation” and “cell cycle control of chromosomal replication” pathways (Fig. 5A). In addition, gene set enrichment analyses showed significant decreases in genes involved in the following KEGG pathways: cell cycle, DNA replication, E2F targets, DNA repair and MYC targets (Supplemental Fig. 5A). RNAseq data from MCF10A cells treated with calcium and ionomycin showed similar results. Ingenuity Pathway analysis demonstrated an increase in the “cell cycle: G2/M DNA damage checkpoint regulation” and “senescence” pathways along with a decrease in “cell cycle control of chromosomal replication” and “cyclins and cell cycle regulation” pathways (Fig. 5B). Gene set enrichment analyses demonstrated a decrease in the expression of genes involved in KEGG hallmark pathways for cell cycle, DNA replication, DNA repair and MYC targets (Supplemental Fig. 5B). Similarly, TGFβ treatment of MCF10A cells mirrored the results of calcium and ionomycin as well as the mammary gland involution pattern. As shown in Fig 5C, Ingenuity Pathway Analysis of RNAseq data from MCF10A cells treated with TGFβ3 showed activation of the “senescence” pathway and inhibition of the “cell cycle control of chromosomal replication” and “cyclins and cell cycle regulation” pathways. Gene set enrichment analyses demonstrated that TGFβ3 inhibited the expression of genes within KEGG hallmark pathways for DNA repair, cell cycle, DNA replication, E2F targets, UV response and MYC targets, V1 and V2 (Supplemental Fig. 5C). Dot-plot showed significantly enriched cell cycle related transcription factors and pathways (Fig. 5D). RNAseq based DoRothEA (Discriminant Regulon Expression Analysis) analysis confirmed that the activity of transcription factors involved in cell cycle progression was downregulated in MCF10A cells treated with either high calcium or TGFβ3 (Fig. 5E). These changes support the idea that mammary gland involution, elevations in intracellular calcium and treatment with TGFβ3 all are associated with inhibition of cell cycle progression.

**Figure 5.**
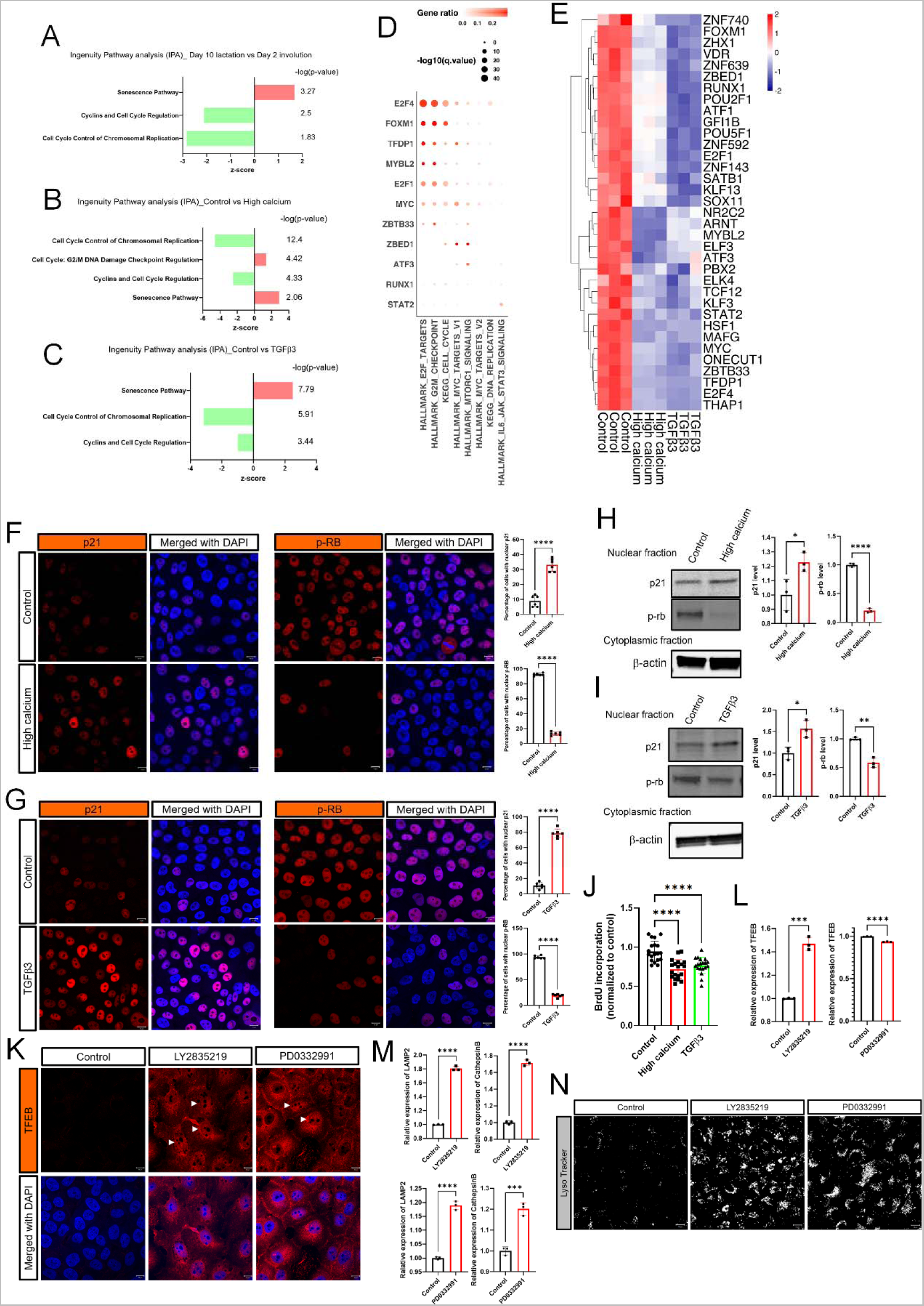
Intracellular calcium and TGFβ3 increase TFEB through inhibition of cell cycle progression. A-C) Ingenuity Pathways Analysis (IPA) highlighting cell cycle related pathways from RNAseq results comparing day 10 lactation with day 2 involution in the mammary gland (A), control and high calcium treated- (B), and control and TGFβ3 treated- MCF10A cells (C). D) Dotplot of significantly enriched pathways related to cell cycle. E) DoRothEA (Discriminant Regulon Expression Analysis) analysis compared control to either high calcium condition or TGFβ3-treated MCF10A cells. F) Immunofluorescence for p21 and p-RB under high calcium conditions. G) Immunofluorescence for p21 and p-RB in MCF10A cells exposed to TGFβ3. H) Western blot analysis of nuclear p21 and p-RB in MCF10A cells under high calcium conditions (n=3). I) Western blot analysis of nuclear p21 and p-RB in MCF10A cells exposed to TGFβ3. (n=3). J) Measurement of BrdU incorporation in MCF10A cells at control versus high calcium conditions, or after treatment with TGFβ3. K) Immunofluorescence for TFEB in MCF10A cells exposed to 2.5μM LY2835219 or 5μM PD0332991. L) TFEB mRNA expression assessed by QPCR in MCF10A cells exposed to LY2835219 or PD0332991. (n=3). M) LAMP2 and Cathepsin B mRNA expression assessed by QPCR in MCF10A cells exposed to LY2835219 or PD0332991. (n=3). N) LysoTracker staining of MCF10A cells at baseline, and after treatment with LY2835219 or with PD0332991. All scale bars represent 10μm.

TGFβ signaling is known to cause cell-cycle arrest in the G1 phase, in part, by increasing the expression of the CDK inhibitor p21, which inhibits CDK4/6 activity and results in hypophosphorylation of retinoblastoma protein Rb (52–54). We therefore examined p21 expression and phosphorylation of Rb in MCF10A cells treated with either calcium/ionomycin or TGFβ3, In both instances, we saw an increase in the proportion of cells with nuclear staining for p21 and a decrease in the proportion of cells staining for nuclear pRb (Fig. 5F&G). We confirmed these immunofluorescence patterns by also performing immunoblots for p21 and pRb in nuclear extracts from MCF10A cells treated with either calcium/ionomycin or TGFβ3. As shown, both calcium/ionomycin and TGFβ3 caused an increase in p21 levels and a decrease in pRb levels in nuclear extracts of MCF10A cells (Fig. 5H&I). As predicted by these results, calcium/ionomycin and TGFβ3 reduced BrdU (Bromodeoxyuridine / 5-bromo-2’- deoxyuridine) incorporation as well (Fig. 5J). Finally, we used two CDK4/6 inhibitors, LY2835219 (Abemaciclib) and PD0332991 (Palbociclib), both of which are used in breast cancer patients, to see whether induction of cell cycle arrest in MCF10A cells would reproduce the effects of calcium/ionomycin or TGFβ3 on TFEB activity. Treatment of MCF10A cells with either PD0332991 or LY2835219 led to an increase in total and nuclear immunofluorescence for TFEB (Fig. 5K). Interestingly, LY2835219, but not PD0332991, also increased TFEB gene expression (Fig. 5L). Despite this difference, gene expression was increased for LAMP2 and Cathepsin B, both known targets of TFEB (Fig. 5M). Both treatments also increased lysosomal mass (Fig. 5N). Since we had previously shown that TGFβ treatment led to a reduction in SOCS3 protein levels and increase in gene expression, we also examined the effects of LY2835219 and PD0332991 on SOCS3 protein levels and *Socs3* gene expression. LY2835219 and PD0332991 treatment led to the same reciprocal decrease in SOCS3 protein levels and increase in *Socs3* gene expression as we had seen with TGFβ3 treatment (Supplemental Fig. 6A&B). The effect of LY2835219 and PD0332991 was not blunted by a pan-TGFβ blocking antibody (Supplemental Fig. 6A&B). Taken together, these data suggest that, during early involution, increased intracellular calcium and TGFβ3 act to induce cell cycle arrest, which, in turn, increases TFEB expression, nuclear localization and signaling to increase lysosome biogenesis, part of a coordinated LDCD pathway (Supplemental Fig. 6C).

## Discussion

The coordinated death of MECs upon weaning is an important feature of mammalian reproduction that allows for the cyclical production of milk following multiple pregnancies, while avoiding the energetic burden of maintaining constant milk production between pregnancies (1, 3, 4, 7). Therefore, it is important to better understand the molecular mechanisms that underlie this process. A series of studies have provided the following working model of early involution: alveolar distension due to milk retention increases cytokine production and activates STAT3, which, in turn, triggers lysosome-mediated cell death pathways (2, 7, 8, 10, 13, 14, 16, 24, 28, 29). However, the mechanisms by which milk stasis activates cytokine production and STAT3 phosphorylation have been less clear. We now present evidence demonstrating that milk stasis rapidly decreases the expression of the calcium pump, PMCA2, causing a dramatic and sustained increase in intracellular Ca^2+^ concentration. This rise in intracellular Ca^2+^ activates LIF, TGFβ3, and IL-6 production, inhibits cell cycle progression, triggers SOCS3 degradation, activates STAT3 signaling, and activates TFEB signaling, all of which contribute to the initiation of LDCD. Interestingly, the fall in PMCA2 levels as well as the increase in intracellular Ca^2+^ generally precede, and are sufficient for the increase in LIF, TGFβ3 and IL6 expression, all cytokines that have previously been suggested to mediate the effects of milk stasis on cell death (8-10, 13, 14). In addition, increased cellular Ca^2+^ levels correlate with an inhibition in cell cycle progression and increased TFEB signaling. Finally, the effects of calcium appear to be mediated, at least in part, by increased TGFβ signaling. Our observations suggest a working model (Supplemental Fig. 6C), whereby the rise in intracellular Ca^2+^ acts as an early biochemical signal for activation of STAT3. Increased intracellular calcium also increases local cytokine production, TFEB activation and lysosome biogenesis leading to an amplification of STAT3 activation, and an increase in lysosomal mass, both of which support the full development of LDCD.

Our results demonstrate that increased intracellular Ca^2+^ activates STAT3 by decreasing SOCS3 levels. The decrease in SOCS3 protein occurs despite a reciprocal increase in SOCS3 mRNA levels, suggesting that increased Ca^2+^ triggers degradation of SOCS3 protein, disinhibiting STAT3-induced expression of the *Socs3* gene. Sutherland and colleagues had previously reported that mammary-specific disruption of the *Socs3* gene led to premature STAT3 phosphorylation and cell death during lactation, mirroring the phenotype of lactating PMCA2-null mice (33, 34). Furthermore, the authors demonstrated an increase in *Socs3* mRNA in control animals 24-hours after weaning, findings consistent with our data. Additionally, we found that the calcium-induced reduction in SOCS3 levels can be mediated by increased TGFβ signaling and that the effects of TGFβ3 on SOCS3 appear to involve the inhibition of cell cycle progression as the reductions in SOCS3 levels can be reproduced by pharmacologic inhibition of CDK4/6 activity. SOCS3 has previously been shown to interact with cell cycle regulators but it is thought to contribute to cell cycle arrest by magnifying p53- mediated upregulation of the cyclin dependent kinase inhibitor, CDKN1A or p21 (55, 56). Our observations of decreased levels of SOCS3 in the setting of increased p21 levels do not fit this model; therefore, further work will be required to understand how inhibition of CDK4/6 decreases SOCS3 protein levels in mammary epithelial cells. Some of this effect may be mediated by induction of TFEB (Fig. 4), although understanding this connection will also require additional study.

Activation of the LDCD pathway during early mammary gland involution is associated with an increase in the number and size of lysosomes. Lysosome biogenesis is coordinated by the transcription factor, TFEB, which upregulates a network of genes encoding proteins important to the structure and function of lysosomes (42, 43, 57). We found that milk stasis and loss of PMCA2 expression both upregulate TFEB expression and nuclear localization *in vivo*. This was associated with an increase in TFEB-regulated genes (44). We also demonstrated that increased intracellular Ca^2+^ increases TFEB expression and nuclear localization in mammary epithelial cells *in vitro* and this was also associated with an upregulation of TFEB-regulated genes based on RNAseq data. Furthermore, as with SOCS3 degradation, the activation of TFEB in response to calcium appears to be mediated, in part, by TGFβ signaling since TGFβ3 can reproduce the effects of increased intracellular calcium. Furthermore, our data demonstrate a clear reciprocal relationship between activation of TFEB pathways and inhibition of cell cycle progression based on RNAseq data in involuting mammary glands *in vivo* and in MCF10A cells treated with either calcium/ionomycin or TGFβ3. These data suggest that the inhibition of cell cycle progression may mediate the effects of intracellular calcium and TGFβ3 on TFEB expression and nuclear localization. Although intracellular calcium and TGFβ have complex relationships with cell cycle regulation (51), inhibition of cell cycle progression has been shown to promote nuclear localization of TFEB and activation of TFEB-associated gene expression (45, 46). It has been shown that CDK4/6 can directly phosphorylate TFEB leading to its nuclear exclusion and degradation (45), findings consistent with our observations that pharmacologic inhibition of cell cycle progression using 2 different CDK4/6 inhibitors increases total and nuclear TFEB levels as well as the expression of the downstream TFEB target genes cathepsin B and LAMP2. Thus, we propose that intracellular calcium and TGFβ3 increase TFEB signaling by inhibiting CDK-mediated cell cycle progression.

Although not all secretory epithelial cells die during the first phase of involution, it appears that a majority of cells demonstrate increased GCaMP6f fluorescence by 24- hours after teat sealing (Fig. 1C). Interestingly, there is more heterogeneity in GCaMP4f fluorescence 24-hours after weaning as suggested in Fig. 2E. This may suggest variability in either the degree of intracellular calcium overload and/or the response to elevated calcium among epithelial cells. It is interesting to speculate whether this might dictate which cells survive the first phase of involution. However, it is difficult to co-register calcium accumulation in vivo with activation of LDCD using intravital calcium imaging and our models in vitro demonstrate uniform elevations in intracellular calcium (Supplemental Figure 2A). Further work will be needed to address potential heterogeneity in these responses.

In summary, we present data identifying intracellular calcium as an important early signal linking milk stasis to mammary epithelial cell death. We found that milk retention causes a decrease in PMCA2 expression and a rise in intracellular Ca^2+^. The increase in intracellular Ca^2+^, in turn, activates STAT3 by degrading SOCS3, which is mediated by alterations in TGFβ signaling and cell cycle arrest. These same pathways also mediate the effects of elevated intracellular calcium on TFEB signaling and lysosome expansion after weaning. We propose that these events function as key proximal signals initiating and amplifying LDCD in MECs during early involution, a process that triggers the death of secretory MECs and, ultimately, initiates the preparation of the mammary gland for a new cycle of reproduction.

## Materials and Methods

### Cell culture

MCF10A cells were grown in 2 dimensional monolayer on plastic, and were cultured in DMEM/F12 (Gibco-Life Technologies) containing 5% horse serum, EGF (100ng/ml), hydrocortisone (1mg/ml), cholera toxin (1mg/ml), insulin (10mg/ml), and pen/strep (Gibco-Life Technologies) at 37°C in 5% CO2 (58). In some experiment, cells were cultured under high calcium conditions (10 mM calcium + 1 μM ionomycin) for 16 hrs, or treated with TGFβ3 (10 nM) for 16 hrs. To inhibit STAT3, cells were treated with Galiellaclactone (10 μM) (Tocris Bioscience, Atlanta, GA) for 16 hrs. To inhibit cell cycle progression, cells were treated with LY2835219 (2.5μM) and PD0332991 (5μM) (Selleckchem, Randnor, PA) for 16 hrs.

### Genetically-altered mice

PMCA2wt/dfw-2J mice were obtained from Jackson Laboratory (CByJ.A- Atp2b2dfw-2J/J, stock number 002894). Ai95(RCL-GCaMP6f)-D (Ai95) were a gift of the Lawrence Cohen laboratory at Yale University and were crossed with BLG-Cre mice, which were the gift of the Christine Watson Laboratory at the University of Cambridge. All animal experiments were approved by the Yale Institutional Animal Care and Use Committee.

### Immunofluorescence

Cells were grown on coverslips, fixed in 4% paraformaldehyde for 20 min, permeabilized with 0.2 % Triton X100 for 10 mins, washed 3 times with PBS and incubated with primary antibody overnight at 4°C. The cells were washed 3 times with PBS and incubated with secondary antibody for 1 hour at room temperature. After washing, coverslips were mounted using Prolong Gold antifade reagent with DAPI (Invitrogen). Paraffin-embedded tissue sections were cleared with histoclear (National Diagnostics) and graded alcohol using standard techniques. Antigen retrieval was performed using 7mM citrate buffer, pH 6.0 under pressure. Sections were incubated with primary antibody overnight at 4°C and with secondary antibody for 1 hour at room temperature. Coverslips were mounted using Prolong Gold antifade reagent with DAPI (Invitrogen). All images were obtained using a Zeiss 780 confocal microscope and Zeiss LSM 880, and settings were adjusted to allow for detection of fine membrane structure. Primary antibodies were against: LAMP2 (ab13524) from Abcam (Cambridge, MA); PMCA2 (PA1-915) from Thermo Scientific (Waltham, MA); cathepsin B (PA5-17007) and TFEB (PA5-96632) from Invitrogen (Grand Island, NY); Phospho- STAT3 (9145), p21 (2947), p-Rb (8516) and NFAT (5861) from cell signaling (Danvers, MA).

### Immunohistochemistry

Paraffin-embedded tissue sections were cleared with histoclear (National Diagnostics) and graded alcohol using standard techniques. Immunohistochemistry was performed using standard techniques (59). Antigen retrieval was accomplished by heating sections in 7 mM or 10 mM citrate, under pressure. Sections were incubated with primary antibody overnight at 4°C. Staining was detected using Vector Elite ABC Kits (Vector Laboratories, Burlingame, CA, USA) and 3,3-diaminobenzidine as chromogen (Vector Laboratories). Primary antibodies were against: phospho-STAT3 (9145), phospho-STAT5 (9314), CREB (9197), phospho-CREB (9198), and NFAT (5861), all from cell signaling (Danvers, MA) as well as TFEB (PA5-96632) from Invitrogen (Grand Island, NY).

### Intravital multiphoton microscopy

Mice were initially anaesthetized with an intraperitoneal injection of ketamine (15mg/mL) and xylazine (1mg/mL) in PBS and maintained throughout the course of the experiment with vaporized isoflurane, 1.5% in oxygen, on a heating pad maintaining temperature at 37°C. The abdomen was shaved using a mechanical trimmer and depilatory cream and the inguinal mammary gland was surgically exposed on a skin flap. The surrounding tissue was pinned to a silicone mount to stabilize and prewarmed PBS (37°C) was applied topically to the flap throughout the imaging procedure. A coverslip mounted on a micromanipulator was lowered onto the mammary gland prior to imaging.

Image stacks were acquired with a LaVision TriM Scope II (LaVision Biotec) microscope equipped with a Chameleon Vision II (Coherent) multiphoton laser. The laser was tuned to 880nm, focused through a ×20 water immersion lens (N.A. 1.0; Olympus) and scanned a field of view of 0.5 mm^2^ at 800 Hz (0.48µm/pixel). Serial optical sections were acquired in 3-μm steps to image a total depth of ∼70μm of tissue. Larger regions were visualized using a motorized stage to automatically acquire sequential fields of view in a 3x3 grid with 4% overlap between regions. Laser power and imaging settings was consistently maintained between all replicates. Emitted fluorescence was collected through two non-descanned detectors and separated through a dichroic (490nm) and bandpass filters (435/90 = blue, 525/50 = green).

Image stacks were initially stitched by a grid/collection stitching plugin in Fiji before importing into Imaris software v9.2.1 (Bitplane) for three-dimensional volume rendering. Surfaces were created based on the green-fluorescent signal and manually segmented into individual alveoli for analysis of their mean fluorescent intensity.

In vivo results represent samples from 3 individual mice for teat sealing experiments and 2 mice for the pup withdrawal and reintroduction experiments. An unpaired Student’s *t*-test was used for all analyses with a *P* value of less than 0.05 accepted as indicating a significant difference. Statistical calculations were performed using the Prism (GraphPad).

### Immunoblotting

Protein extracts were prepared using standard methods (58, 60), subjected to SDS-PAGE and transferred to a nitrocellulose membrane by wet western blot transfer (Bio-Rad). The membrane was blocked in TBST buffer (TBS + 1% Tween) containing 5% milk for 1 hour at room temperature. The blocked membranes were incubated overnight at 4 °C with primary antibodies in Odyssey blocking buffer, 927-40000, washed 3 times with TBST buffer, and then incubated with secondary antibodies provided by LI-COR for 2 hours at room temperature. After 3 washes with TBST buffer, the membranes were analyzed using the ODYSSEY Infrared Imaging system (LI-COR). Primary antibodies were against: PMCA2 (PA1-915) from Thermo Scientific (Waltham, MA); phospho- STAT3 (9145), STAT3 (9139), p21 (2947), p-Rb (8516), SOCS3 (2923), Phospho-S6 Ribosomal Protein (Ser235/236) (4858), S6 Ribosomal Protein (5G10) (2217), Phospho-p70 S6 Kinase (Thr389) (9205), p70 S6 Kinase Antibody (9202), Phospho- ULK1 (Ser757) (D7O6U) (14202), and ULK1 (D8H5) (8054) from cell signaling (Danvers, MA); SOCS3 (HPA068569) from sigma (Burlington, MA); mouse (sc-69879) and rabbit (sc-130656) β-actin from Santa Cruz (Dallas, TX); TFEB (PA5-96632) and cathepsin B (PA5-17007) from Invitrogen (Grand Island, NY); SOCS3 (ab16030) from Abcam (Cambridge, MA). All immunoblot experiments were performed at least 3 times and representative blots are shown in the figures.

### Knockdown Cell Line

A stable cell line expressing shRNA directed against TGFBR2 and TFEB were generated by transducing cells with commercially prepared lentiviruses containing three individual shRNA directed against TGFBR2 (sc-36657-v) and TFEB (sc-38509-v) mRNA (Santa Cruz). Cells were cultured in 6-well plates and infected by adding the shRNA lentiviral particles to the culture for 48 hours per the manufacturer’s instructions. Stable clones expressing the specific shRNAs were selected using 5μg/ml of puromycin (Gibco-life technologies) and pooled to generate the cells used in the experiments.

### Cell Transfections

Constructs encoding pCAG cyto-RCaMP1h (plasmid #105014), and GcaMP3- TRPML were a gift of Haoxing Xu in the University of Michigan. SKBR3 cells were transfected using Fugene6 transfection reagent (Invitrogen) according to the manufacturer’s instructions.

### RNA Extraction and Real-Time RT-PCR

RNA was isolated using TRIzol (Invitrogen). Quantitative RT-PCR was performed with the SuperScript III Platinum One-Step qRT-PCR Kit (Invitrogen) using a Step One Plus Real-Time PCR System (Applied Biosystems) and the following TaqMan primer sets: human and mouse Lif (Hs01055668_m1 and Mm00434761_m1), mouse PMCA2 (Mm00437640_m1), human and mouse CD14 (Hs02621496_s1 and Mm00438094_g1), human and mouse LBP (Hs01084628_m1 and Mm00493139_m1), human and mouse IL6 (Hs00174131_m1 and Mm00446190_m1), human and mouse SOCS3 (Hs02330328_s1 and Mm00545913_s1), human and mouse TGFβ3 (Hs01086000_m1 and Mm00436960_m1), human LAMP2 (Hs00174474_m1), human ATG7 (Hs00893766_m1), and human CTSB (Hs00947439_m1), and human and mouse TFEB (Hs00292981_m1 and Mm00448968_m1). Human HPRT1 (4325801) and mouse GAPD (4351309) were used as reference genes (Invitrogen). Relative mRNA expression was determined using the Step One Software v2.2.2 (Applied Biosystems).

### Bulk RNA sequencing

RNA sequencing was performed by the Yale Center for Genome Analysis using the Illumina NovaSeq 6000 system, with 2x100 bp paired end. The sequencing reads were aligned onto the mouse GRCm38/mm10 and the Human GRCh38/hg38 reference genomes using the HISAT2 ver.2.1.0 (61) software. The mapped reads were converted into the count matrix using StringTie2 ver. 2.1.4 (62) with the default parameters, and provided to DESeq2 ver. 1.32.0 (63) to identify differentially expressed genes (DEGs) based on a negative binomial generalized linear models. Genes that satisfy |Log2 Fold Change| ≥ 0.25 and adjusted p-values < 0.05 were considered as statistically significant. The data visualization of the DEGs along with TFEB related genes and hierarchical clustered heatmaps were performed using the EnhancedHeatmap package 1. (64) in R.

### Gene regulatory network analysis

Regulons were defined by a gene regulatory network called DoRothEA (65) containing a collection of TF - target gene interactions. decoupleR package was used to estimate regulon activities by a multivariate linear model (mlm) from the transcriptome data. Enrichment analysis of each regulon for HALLMARK and KEGG gene sets was performed by Fisher’s exact test utilizing clusterProfiler (66) package.

### Statistics

Statistical analyses were performed with Prism 7.0 (GraphPad Software, La Jolla, CA). Statistical significance was determined by using unpaired t test for comparisons between 2 groups and one-way ANOVA for groups of 3 or more. All bar graphs represent the mean±SEM, * denotes p<0.05, ** denotes p<0.005, *** denotes p<0.0005, **** denotes p<0.00005.

## Data availability

All data and information are included in the article and/or the SI appendix. RNA-seq data was deposited in NCBI’s GEO (Accession number GSE221243).

## Acknowledgements

We thank the Lawrence Cohen laboratory at Yale University for advice on genetically encoded calcium sensor mice and the gift of the Ai95(RCL-GCaMP6f)-D (Ai95) mice for our experiments. We thank Dr. Christine Watson from Cambridge University for valuable conversations and advice as well as the gift of the BLG-Cre mice. We thank Dr. Haoxing Xu in the University of Michigan for the plasmid encoding bGcaMP3- TRPML. We thank the Yale Center for Advanced Light Microscopy Facility for their assistant, and the Bruker Opterra Swept Filed Microscope was funded by shared instrument grant # NIH S10 OD023598. Our experiments were supported by the following grants: NRF-2018R1C1B6002803 (W. Kim) and NRF-2022R1A4A2000827(J. Choi) from National Research Foundation of Korea, RO1 GM105718 from NIH to S. Ferguson, and R01 HD100468 and R01 HD076248 from the NIH to J. Wysolmerski.

## Conflict of interest

Nothing declared.

## Notes

### Competing Interest Statement

The authors have declared no competing interest.

### Summary of Updates

Figure 3&4 revised

