## Supplemental information for "Intracellular Calcium links Milk Stasis to Lysosome Dependent Cell Death by Activating a TGFβ3/TFEB/STAT3 Pathway Early during Mammary Gland Involution"

Supplemental Figure_1

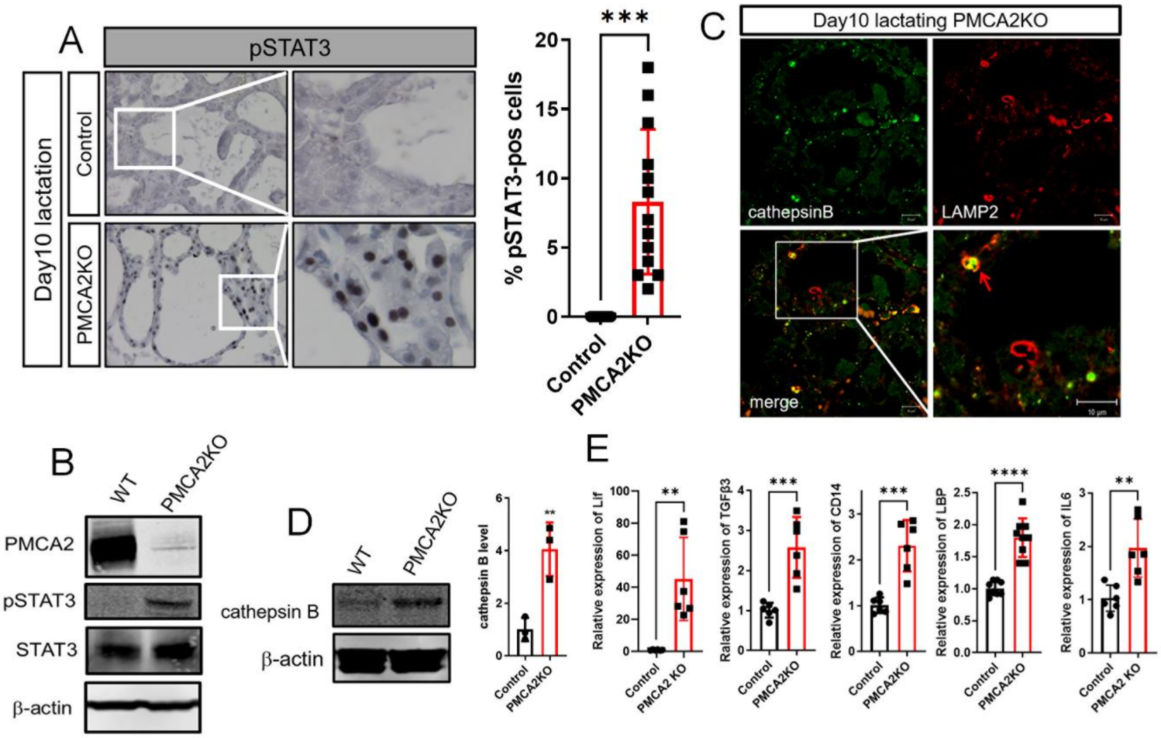

**Supplemental Figure 1. Loss of PMCA2 prematurely activates LDCD during lactation.** A) Immunohistochemistry for pSTAT3 in mammary glands of control and PMCA2-null mice on day 10 of lactation. (n=3). B) Western blot analysis of PMCA2, pSTAT3, and STAT3 in mammary glands from control and PMCA2 KO mice on day 10 of lactation. (n=3). C) Immunofluorescence for Cathepsin B and LAMP2 in mammary glands from control and PMCA2 KO mice on day 10 of lactation. Red arrow; enlarged lysosome containing cathepsin B. D) Western blot analysis for Cathepsin B in mammary gland lysates from control and PMCA2 KO mice on day 10 of lactation (n=3). E) Lif, TGFβ3, CD14, LBP, and IL6 mRNA expression assessed by QPCR in mammary glands from control and PMCA2 KO mice on day 10 of lactation. (n=6). All scale bars represent 10μm.

Supplemental Figure_2

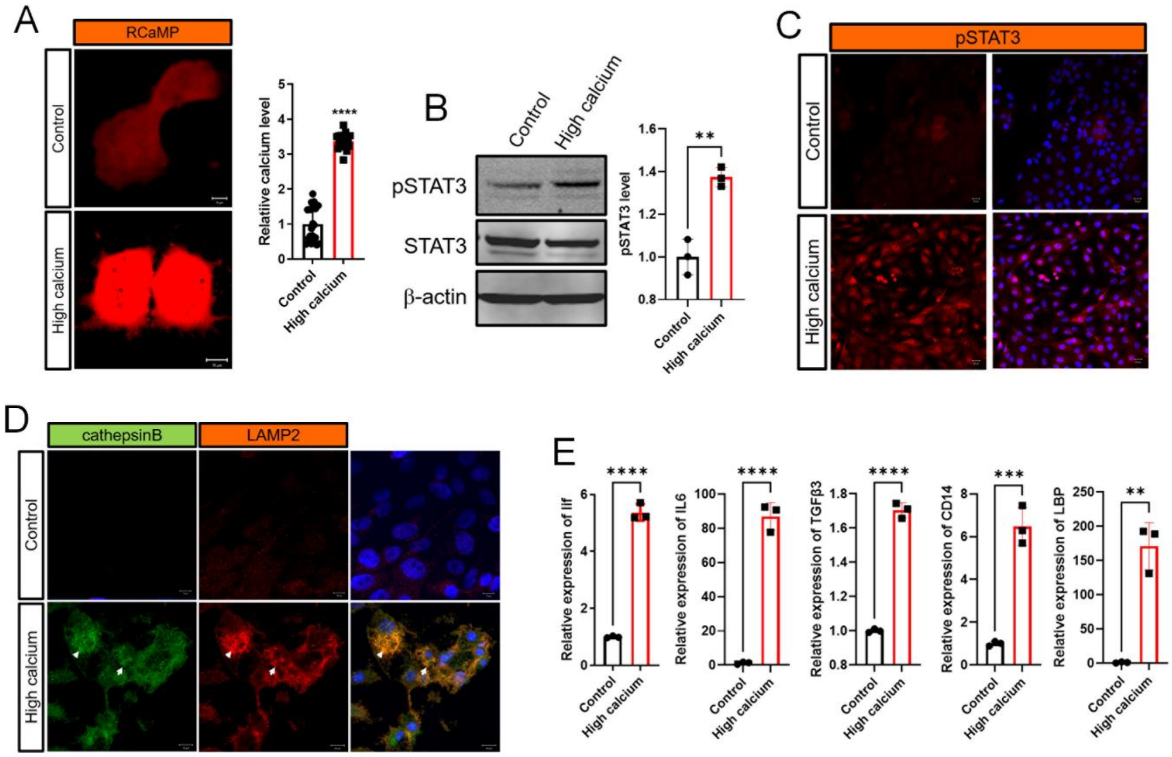

**Supplemental Figure 2. Increased Intracellular Calcium activates STAT3 *in vitro*.** A) Live cell imaging of MCF10A cells expressing the RCaMP cytoplasmic calcium indicator in response to control media (top) and treatment with 10mM calcium + 1μM ionomycin. n= 20 cells for each of 3 experiments. B) Western analysis of total STAT3 and pSTAT3 from MCF10A cells under control and high calcium (10mM calcium + 1μM ionomycin). (n=3) C) Immunofluorescence for pSTAT3 in MCF10A cells under control or high calcium conditions (10mM calcium + 1μM ionomycin). D) Immunofluorescence for LAMP2 and Cathepsin B in MCF10A cells under control or high calcium conditions (10mM calcium + 1μM ionomycin). Scale bars represents 10μm. E) Lif, IL6, TGFβ3, CD14, and LBP mRNA expression in MCF10A cells under control or high calcium conditions (10mM calcium + 1μM ionomycin), as assessed by quantitative RT-PCR (QPCR) (n=3). Bar graphs represent the mean±SEM. ** denotes $p<0.005$, *** denotes $p<0.0005$, **** denotes $p<0.00005$.

Supplemental Figure_3

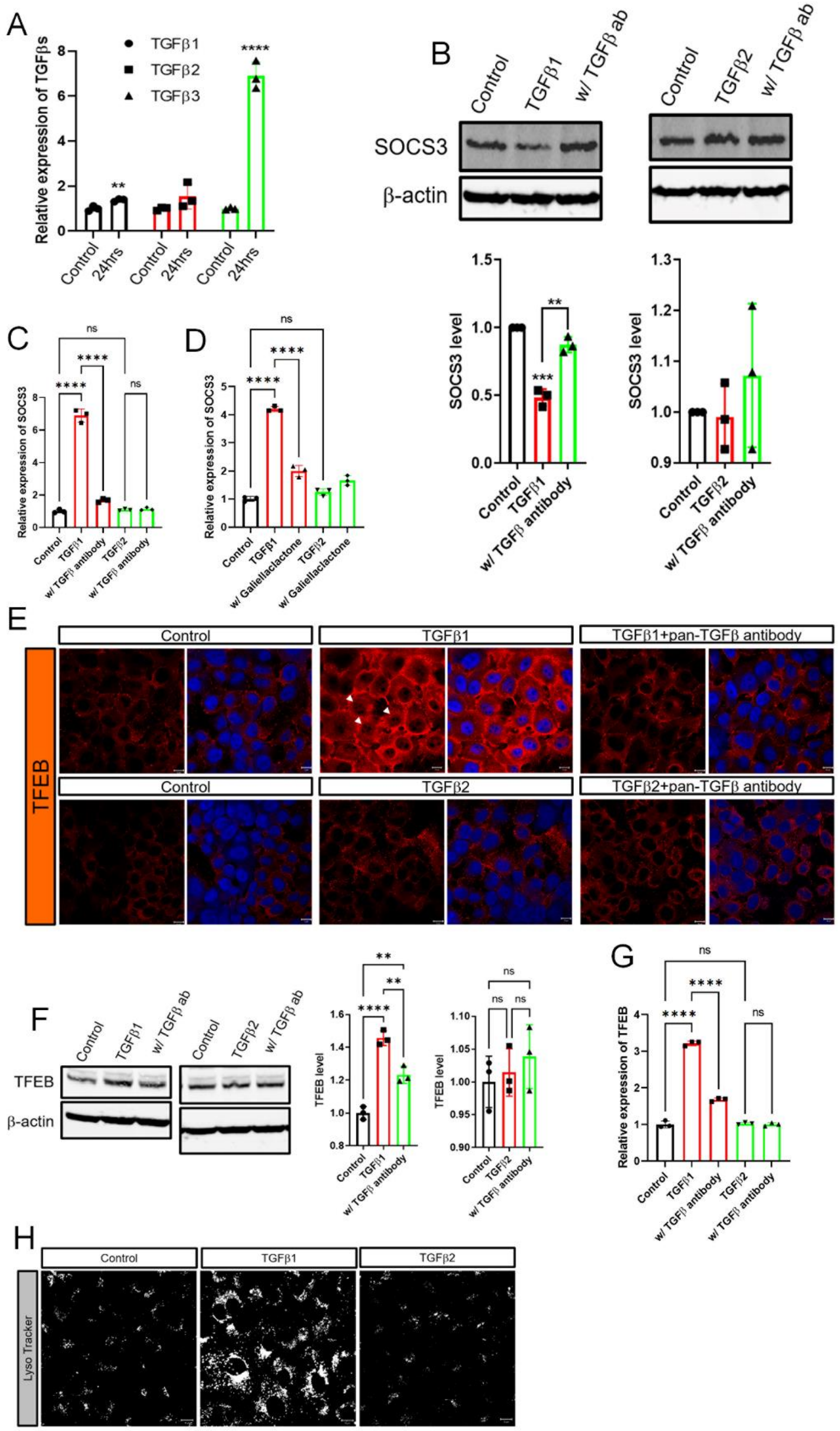


35

**Supplemental Figure 3. TGFβ1 induces SOCS3 degradation and activates TFEB.** A) TGFβ1, 2, and 3 mRNA expression in mammary glands in control and 24 hours post teat-sealing, as assessed by quantitative RT-PCR (QPCR) (n=3). B) Western blot analyses of SOCS3 in MCF10A cells exposed to 10ng/ml TGFβ1 or TGFβ2 with or without co-treatment with 2μg/ml pan-TGFβ antibody. (n=3). C) SOCS3 mRNA expression assessed by QPCR in MCF10A cells exposed to 10ng/ml TGFβ1 or TGFβ2 ± 2μg/ml pan-TGFβ antibody. (n=3). D) SOCS3 mRNA expression assessed by QPCR in MCF10A cells exposed to 10ng/ml TGFβ1 or TGFβ2 ± 10μM/ml of Galiellaclactone (n=3). E) Immunofluorescence for TFEB in MCF10A cells exposed to 10ng/ml TGFβ1 or TGFβ2 ± 2μg/ml pan-TGFβ antibody. Scale bars represents 10μm. F) Western blot analysis of TFEB in MCF10A cells exposed to 10ng/ml TGFβ1 or TGFβ2 ± 2μg/ml pan-TGFβ antibody. (n=3). G) TFEB mRNA expression assessed by QPCR in MCF10A cells exposed to 10ng/ml TGFβ1 or TGFβ2 ± 2μg/ml pan-TGFβ antibody. (n=3). H) LysoTracker staining of MCF10A cells at baseline or after treatment with TGFβ1 or TGFβ2 (10ng/ml). Bar graphs represent the mean±SEM. ** denotes $p<0.005$, *** denotes $p<0.0005$, **** denotes $p<0.00005$.

Supplemental Figure_4

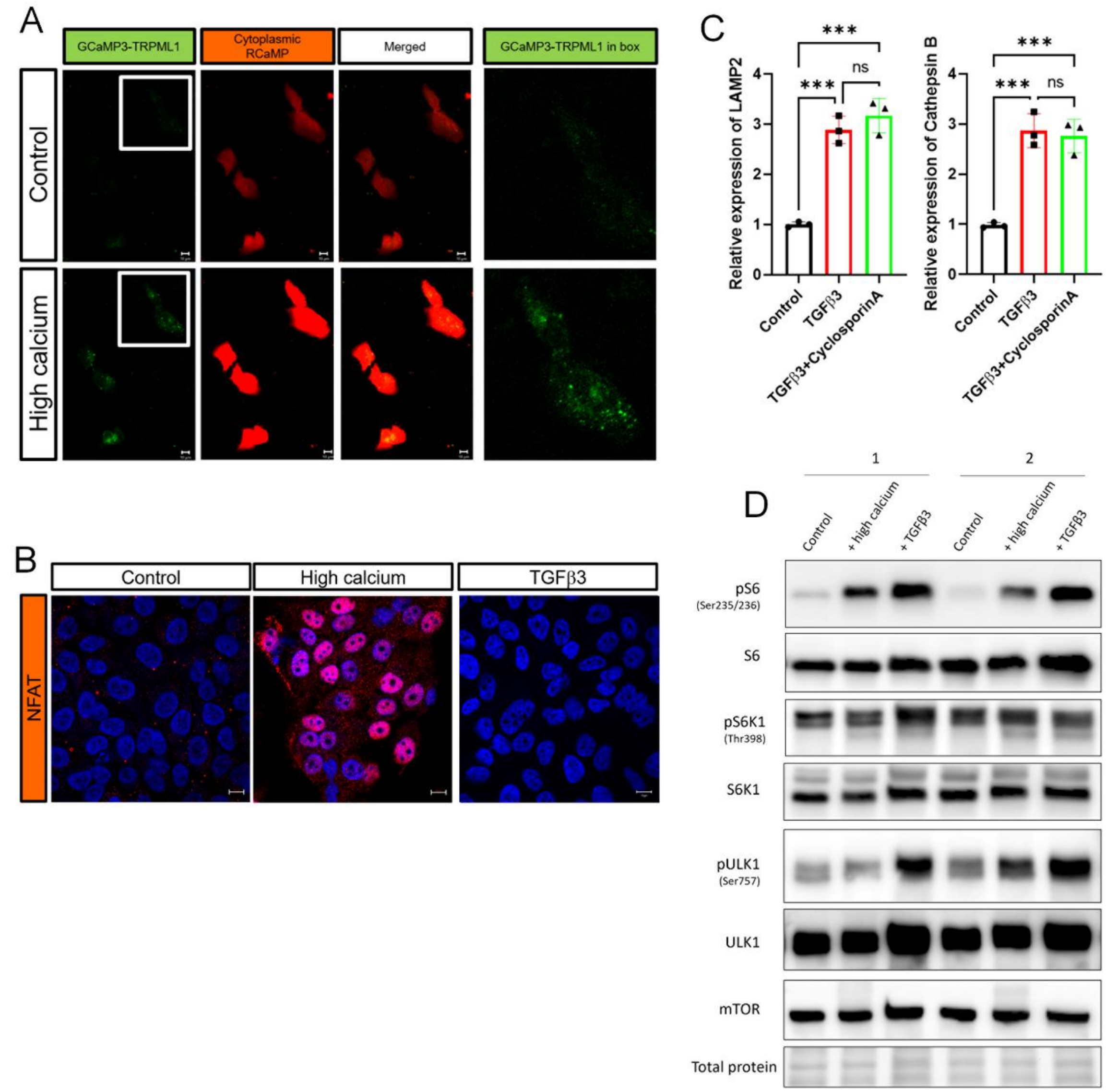

**Supplemental Figure 4. TGFβ3 dependent TFEB activation is not associated with calcineurin and mTOR signaling.** A) Live cell Imaging of GCaMP3-TRPML1 and RCaMP in MCF10A cells grown at control (top) or high calcium (bottom) conditions. Red fluorescence is triggered by cytoplasmic calcium levels. Green fluorescence is triggered by calcium transport out of lysosomes through the TRPML1 calcium pump. B) Immunofluorescence for NFAT in MCF10A cells exposed to high calcium conditions (10mM calcium + 1μM ionomycin) and 10ng/ml TGFβ3. Scale bars represents 10μm. C) LAMP2 and Cathepsin B mRNA expression assessed by QPCR in MCF10A cells exposed to 10ng/ml TGFβ3 ± 1μM/ml Cyclosporin A. (n=3). D) Western blot analysis of pS6 (Ser235/236), S6, pS6K1, S6K1, pULK1 (Ser757), ULK1, and mTOR in MCF10A cells exposed to high calcium conditions (10mM calcium + 1μM ionomycin) and 10ng/ml TGFβ3. Bar graphs represent the mean±SEM. *** denotes $p<0.0005$.

Supplemental Figure_5

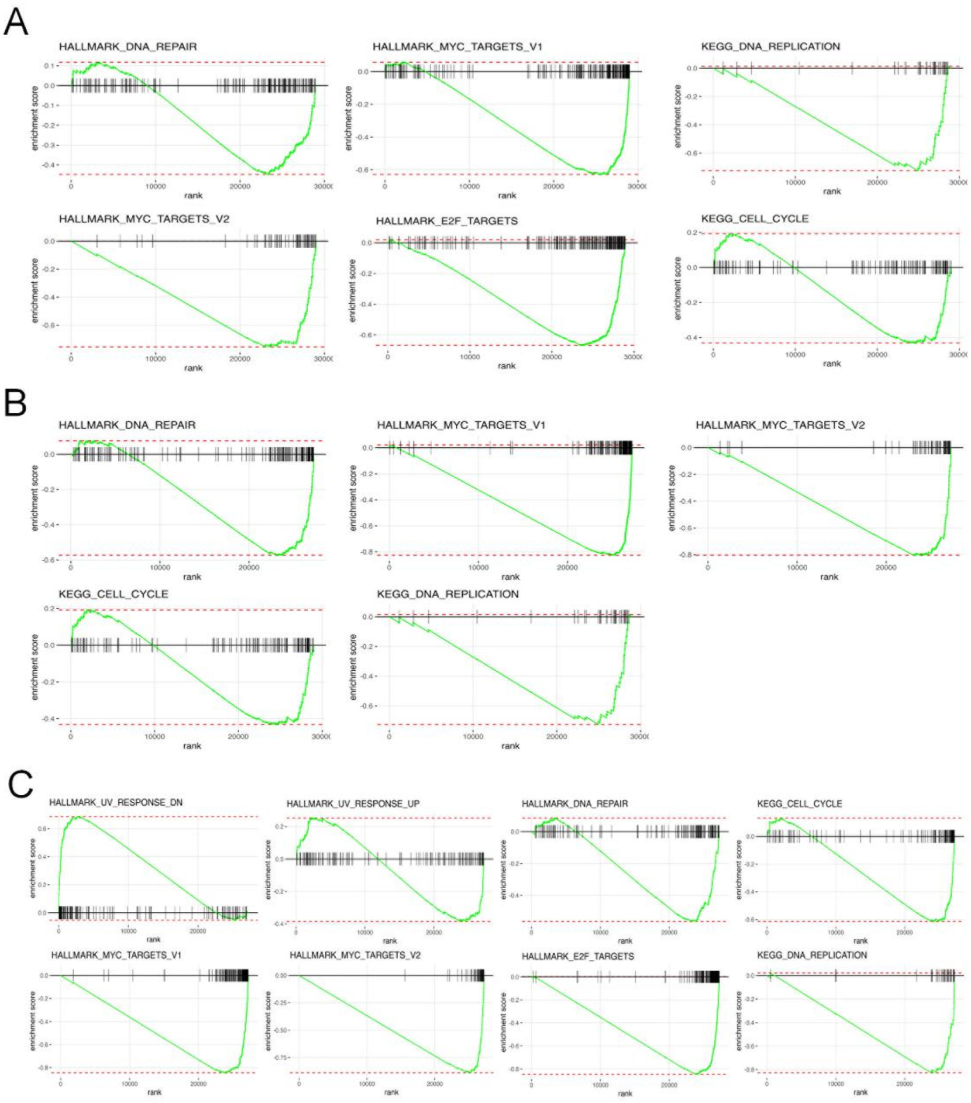

**Supplemental Figure 5. Inhibition of cell cycle progression by high calcium condition and TGFβ3.** A) Hallmark and KEGG gene set enrichment plot of cell cycle related pathways from RNAseq results comparing day 10 lactation with day 2 involution in the mammary gland. B) Hallmark and KEGG gene set enrichment plot of cell cycle related pathways from RNAseq results comparing control and high calcium conditions (10mM calcium + 1μM ionomycin) in MCF10A cells. C) Hallmark and KEGG gene set enrichment plot of cell cycle related pathways from RNAseq results comparing MCF10A cells at baseline and after treatment with 10ng/ml TGFβ3.

Supplemental Figure_6

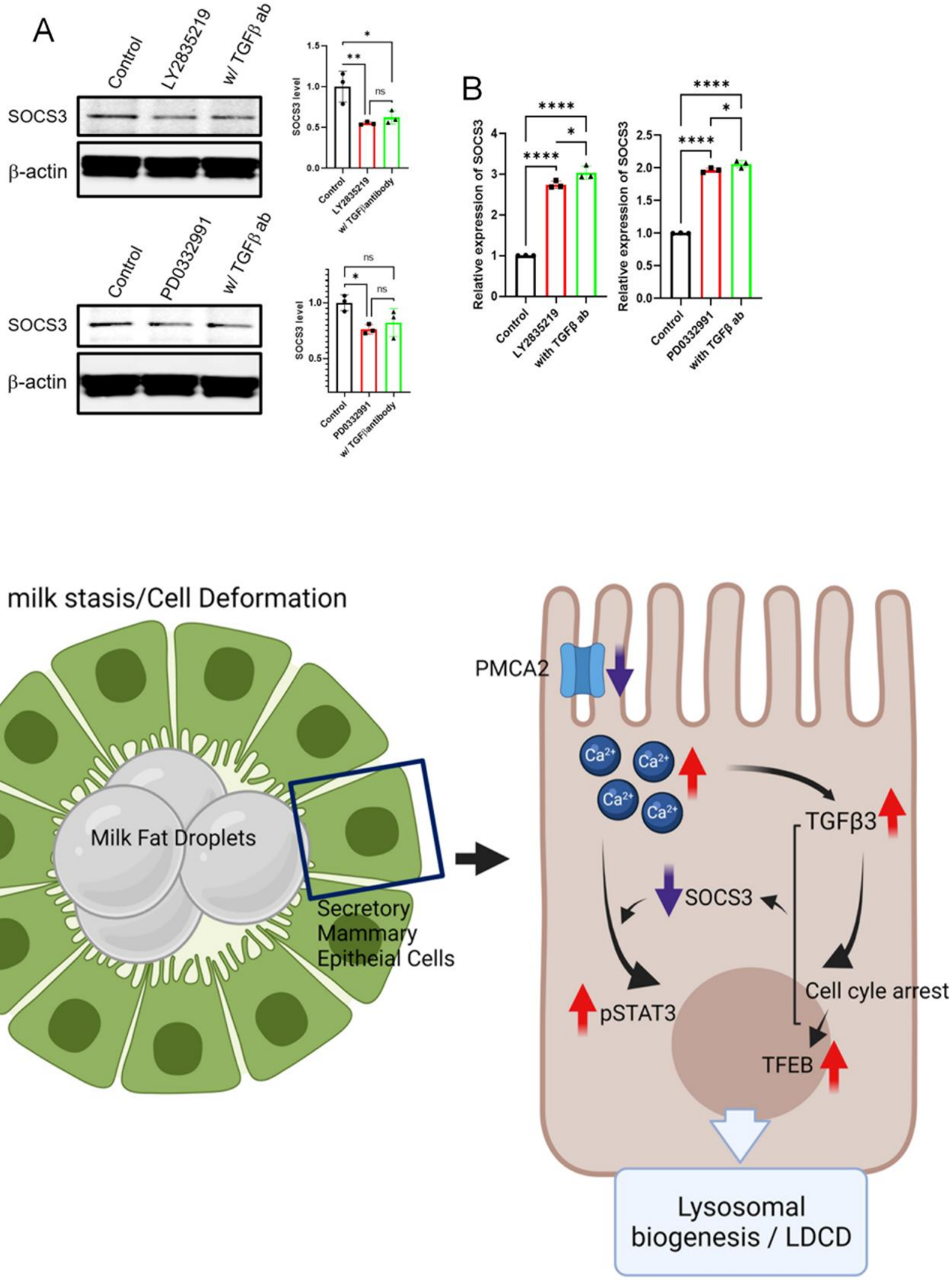

**Supplemental Figure 6. Inhibition of cell cycle progression reduces SOCS3 levels.** A) Western blot analysis of SOCS3 in MCF10A cells exposed to 2.5μM LY2835219 or 5μM PD0332991 ± 2μg/ml pan-TGFβ antibody. (n=3). B) SOCS3 mRNA expression assessed by QPCR in MCF10A cells exposed to 2.5μM LY2835219 or 5μM PD0332991 ± 2μg/ml pan-TGFβ antibody. (n=3). C) Working model illustrating how milk stasis initiates LDCD by decreasing PMCA2 levels and increasing intracellular $Ca^{2+}$. Created with BioRender.com. Bar graphs represent the mean±SEM. * denotes $p<0.05$, ** denotes $p<0.005$, **** denotes $p<0.00005$.

**Supplemental Video 1. Intracellular calcium level in day 10 lactating gland as assessed by GCaMP6f fluorescence. (Control).** Z-stacks of multiphoton laser scanning microscopic images take from control lactating glands from BLG-Cre;Ai95 female mice. Blue fluorescence represents the second harmonic-generated signal from the collagen fibers within the fascia covering the glands. Green fluorescence is derived from increased intracellular calcium in MECs expressing the GCaMP6f calcium indicator.

**Supplemental Video 2. Intracellular calcium level in mammary gland at 4 hours post teat-sealing as assessed by GCaMP6f fluorescence.** Z-stacks of multiphoton laser scanning microscopic images take from 4 hours post teat-sealing from BLG-Cre;Ai95 female mice. Blue fluorescence represents the second harmonic-generated signal from the collagen fibers within the fascia covering the glands. Green fluorescence is derived from increased intracellular calcium in MECs expressing the GCaMP6f calcium indicator.

**Supplemental Video 3. Intracellular calcium level in mammary gland at 8 hours post teat-sealing as assessed by GCaMP6f fluorescence.** Z-stacks of multiphoton laser scanning microscopic images take from 8 hours post teat-sealing from BLG-Cre;Ai95 female mice. Blue fluorescence represents the second harmonic-generated signal from the collagen fibers within the fascia covering the glands. Green fluorescence is derived from increased intracellular calcium in MECs expressing the GCaMP6f calcium indicator.

**Supplemental Video 4. Intracellular calcium level in mammary gland at 24 hours post teat-sealing as assessed by GCaMP6f fluorescence.** Z-stacks of multiphoton laser scanning microscopic images take from 24 hours post teat-sealing from BLG-Cre;Ai95 female mice. Blue fluorescence represents the second harmonic-generated signal from the collagen fibers within the fascia covering the glands. Green fluorescence is derived from increased intracellular calcium in MECs expressing the GCaMP6f calcium indicator.

**Supplemental Video 5. Intracellular calcium level in mammary gland at 24 hours post teat-sealing from BLG-Cre absence Ai95 female mice as assessed by GCaMP6f fluorescence.** Z-stacks of multiphoton laser scanning microscopic images take from 24 hours post teat-sealing from Ai95 female mice (NO Cre). Blue fluorescence represents the second harmonic-generated signal from the collagen fibers within the fascia covering the glands. Green fluorescence is derived from intracellular calcium in MECs expressing the GCaMP6f calcium indicator.

**Supplemental Video 6. Intracellular calcium level in mammary gland at 24 hours after pups were removed as assessed by GCaMP6f fluorescence.** Z-stacks of multiphoton laser scanning microscopic images from 24 hours after pups removed at day 10 of lactation from BLG-Cre;Ai95 female mice. Blue fluorescence represents the second harmonic-generated signal from the collagen fibers within the fascia covering the glands. Green fluorescence is derived from increased intracellular calcium in MECs expressing the GCaMP6f calcium indicator.

**Supplemental Video 7. Intracellular calcium level in mammary gland at 24 hours after pups were reintroduced following 24-hours without suckling as assessed by GCaMP6f fluorescence.** Z-stacks of multiphoton laser scanning microscopic images from 24 hours after pups removed at 24 hours after pups reintroduced following 24-hours without suckling from BLG-Cre;Ai95 female mice. Blue fluorescence represents the second harmonic-generated signal from the collagen fibers within the fascia covering the glands. Green fluorescence is derived from increased intracellular calcium in MECs expressing the GCaMP6f calcium indicator.
